# Analysis of proteins in computational models of synaptic plasticity

**DOI:** 10.1101/254094

**Authors:** Katharina F. Heil, Emilia M. Wysocka, Oksana Sorokina, Jeanette Hellgren Kotaleski, T. Ian Simpson, J. Douglas Armstrong, David C. Sterratt

## Abstract

The desire to explain how synaptic plasticity arises from interactions between ions, proteins and other signalling molecules has propelled the development of biophysical models of molecular pathways in hippocampal, striatal and cerebellar synapses. The experimental data underpinning such models is typically obtained from low-throughput, hypothesis-driven experiments. We used high-throughput proteomic data and bioinformatics datasets to assess the coverage of biophysical models.

To determine which molecules have been modelled, we surveyed biophysical models of synaptic plasticity, identifying which proteins are involved in each model. We were able to map 4.2% of previously reported synaptic proteins to entities in biophysical models. Linking the modelled protein list to Gene Ontology terms shows that modelled proteins are focused on functions such as calmodulin binding, cellular responses to glucagon stimulus, G-alpha signalling and DARPP-32 events.

We cross-linked the set of modelled proteins with sets of genes associated with common neurological diseases. We find some examples of disease-associated molecules that are well represented in models, such as voltage-dependent calcium channel family (*CACNA1C*), dopamine D1 receptor, and glutamate ionotropic NMDA type 2A and 2B receptors. Many other disease-associated genes have not been included in models of synaptic plasticity, for example catechol-O-methyltransferase (*COMT*) and *MAO A*. By incorporating pathway enrichment results, we identify *LAMTOR*, a gene uniquely associated with Schizophrenia, which is closely linked to the MAPK pathway found in some models.

Our analysis provides a map of how molecular pathways underpinning neurological diseases relate to synaptic biophysical models that can in turn be used to explore how these molecular events might bridge scales into cellular processes and beyond. The map illustrates disease areas where biophysical models have good coverage as well as domain gaps that require significant further research.

**Author summary:** The 100 billion neurons in the human brain are connected by a billion trillion structures called synapses. Each synapse contains hundreds of different proteins. Some proteins sense the activity of the neurons connecting the synapse. Depending on what they sense, the proteins in the synapse are rearranged and new proteins are synthesised. This changes how strongly the synapse influences its target neuron, and underlies learning and memory. Scientists build computational models to reason about the complex interactions between proteins. Here we list the proteins that have been included in computational models to date. For good reasons, models do not always specify proteins precisely, so to make the list we had to translate the names used for proteins in models to gene names, which are used to identify proteins. Our translation could be used to label computational models in the future. We found that the list of modelled proteins contains only 4.2% of proteins associated with synapses, suggesting more proteins should be added to models. We used lists of genes associated with neurological diseases to suggest proteins to include in future models.

## Introduction

Activity-dependent synaptic plasticity is necessary for learning and memory [1]. Since the discovery of long term potentiation (LTP) and long term depression (LTD) [2, 3], it has been shown that synaptic plasticity can depend strongly on patterns of pre-and post-synaptic firing [4] and neuromodulators [5]. Forms of plasticity vary between types of synapses and brain region [4], which could be explained by the local proteome, i.e. the expressed proteins and their abundances; PSD-95 knock-outs demonstrate the influence of the proteome on synaptic plasticity [6]. Synaptic plasticity underlies behaviour, as evidenced by the effect of antagonising NMDA receptors [1], and synaptic proteins underlie disease [7].

Synapses have been modelled computationally at various levels of detail. Models at a phenomenological level, such as spike-timing dependent plasticity (STDP) models, link firing patterns in the pre-and postsynaptic neurons to changes in synaptic strength with little or no reference to the underlying molecules [8]. Biophysical models refer to at least some known molecular actors in synaptic plasticity. In 2009 there were at least 117 biophysical postsynaptic signal transduction models [9] and the number is growing [10,11].

Recent advances in tissue and cell extraction techniques and sample processing allow localised proteomes to be determined, e.g. the synapse including the smaller presynaptic or postsynaptic proteomes [12,13]. The most recent analysis of 37 published synaptic proteomic datasets contains 1,867 presynaptic genes, 5,053 postsynaptic genes and 5,862 synaptic genes (with human EntrezID identifiers) respectively. These numbers are large compared to results from individual studies. Nevertheless, data inclusion was highly restrictive and the augmented numbers can be partly explained by higher experimental sensitivity and the broad use of high-throughput techniques (a manuscript containing detailed analysis of the synaptic proteome is in preparation).

These synaptic protein lists make it possible to compare systematically proteins contained in computational models of synapses with those proteins likely to be in the synapse. In this paper we: (1) survey a selection of biophysical models of synaptic plasticity, identifying which proteins are involved in each model, and describing the complexity and detail of description of signalling pathways within the models; (2) compare the proteins in models with synaptic protein lists, thus showing what fraction of synaptic proteins have been considered in models; (3) identify the functional classes of proteins in models; and (4) compare the proteins in models with those involved in neurological diseases. This work should help inform what proteins and pathways should be considered in new modelling efforts. While new datasets offer possibilities for models of greater scope and detail, it is important to understand the foundations that have been laid by existing computational models of synaptic plasticity, which we do thematically before moving to the identification of proteins in models and the discussion of implications of our findings for future synaptic models and model annotation.

## Biophysical models of synaptic plasticity

To set the scene for our analysis of proteins in biophysical models of synapses, we first give an overview of how the questions addressed in models of synaptic plasticity have shaped the development of simulation methods, and describe the main hippocampal, striatal and cerebellar pathways that have been modelled. We categorise simulation methods as non-spatial, spatial or multiscale and as deterministic or stochastic. Table 1 shows examples of simulation packages and associated studies that fall into each category. Rather than using simulators, some studies use bespoke code in languages such as Java, or generic mathematical environments such as MatLab.

**Table 1.**
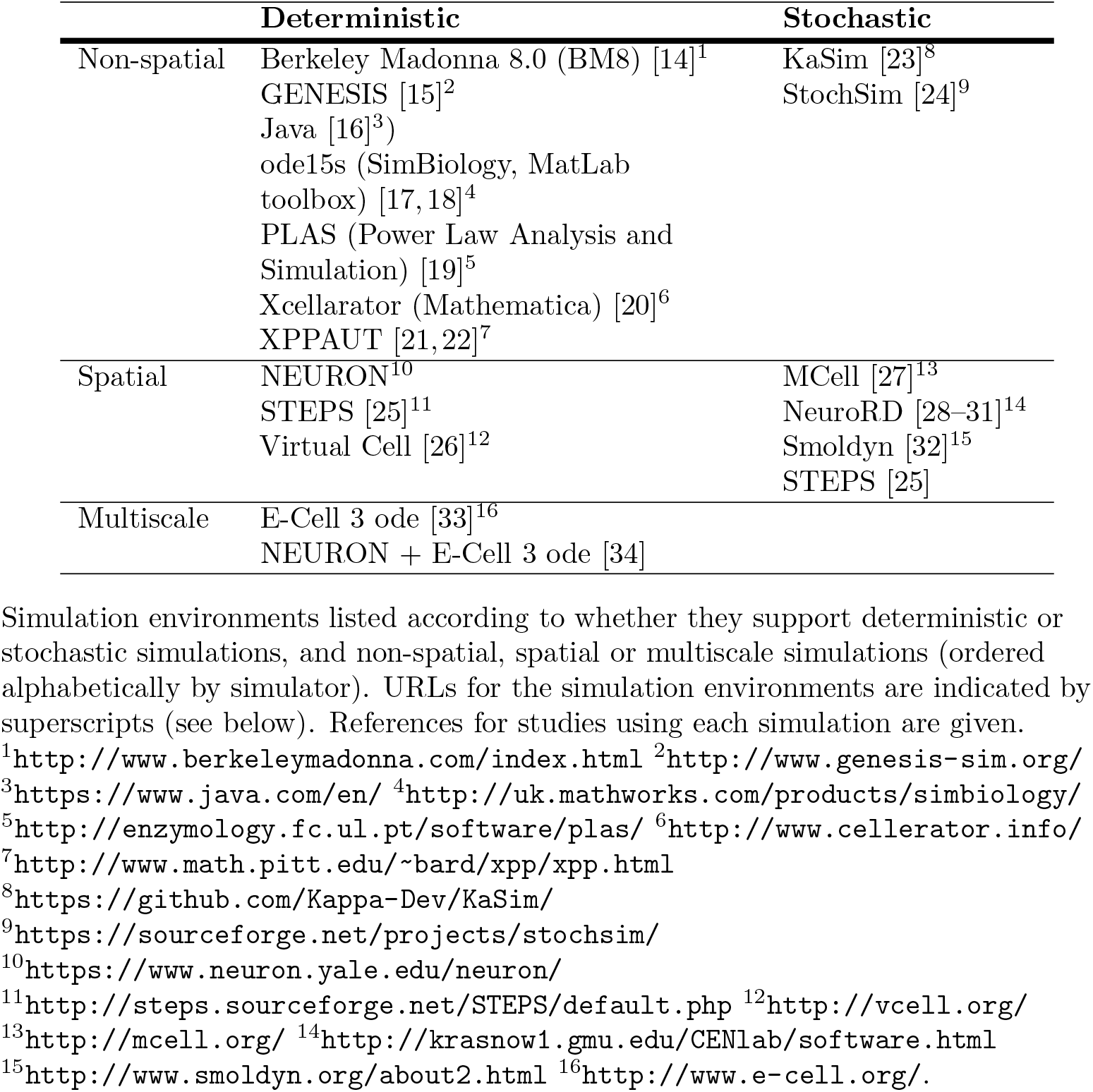
Overview of simulation environments.

### Non-spatial models

Many of the simulation methods and issues associated with models of signalling pathways are found in models of calcium/calmodulin dependent kinase II (CaMKII) and the intricate dynamics of its phosphorylation states and interactions with calcium-bound calmodulin (CaM).

#### Mean field models of CaMKII

In 1985 Lisman [35] advanced the hypothesis, expressed as a mathematical model, that memories could be stored in bistable molecular switches comprised of auto-phosphorylating kinases. Following the discoveries that CaMKII is an autophosphorylating holoenzyme [36] and is a major component of the postsynaptic density (PSD) [37], Lisman and Goldring [38] proposed that CaMKII could form the basis for the auto-phosphorylating switch. Their ordinary differential equations (ODEs) described how the probability of a CaMKII holoenzyme being “on”-the “mean field” - could depend on the calcium concentration and the number of phosphorylation sites required to switch the CaMKII holoenzyme on. Solving these equations demonstrated that the number of CaMKII holoenzymes activated could depend on the duration of the calcium stimulus, thus allowing CaMKII to act as graded rather than binary switch. Furthermore, the time taken for the switch to turn on could be modulated by changing the threshold number of sites that needed to be phosphorylated before the holoenzyme entered an auto-phosphorylated state.

#### Analysis of mean field models

Mean-field ODE models allow stability analysis to be undertaken, which can show, for example, that a model of CaMKII has two stable states - almost fully phosphorylated or almost fully dephosphorylated - within a wide range of calcium concentrations [39]. Stability analysis has also been used to inform how parameters should be set to give a biphasic calcium-synaptic strength curve, with LTD at moderate concentrations of calcium and LTP at high concentrations [22].

#### Stochastic models of CaMKII

In a volume containing *N* reacting molecules of a species, there will be fluctuations of the order of 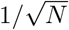 in the concentration of the species predicted by the mean-field solution. For large volumes it follows that stochastic effects can be neglected, but in the ∼1 fl volume of the spine head the number of CaMKII holoenzymes is considerably finite - an average of 30 are seen in electron microscopy (EM) images of immunuogold labelled PSDs [40] - so there will be significant variability between experiments in the same conditions. In order to determine the accuracy of the encoded information for a given number of holoenzymes, Lisman and Goldring [38] used the binomial formula to compute the mean and standard deviation of the number of fully phosphorylated CaMKII holoenzymes, which suggested that graded information could be stored to an accuracy of around 10%.

Rather than deriving variability from mean field simulations, stochastic (“Monte-Carlo”) models can be built. Each run of a stochastic model is generated by drawing random numbers to decide when bonds are made or broken, and when changes in state occur; the variability of the model is obtained by analysing multiple runs. A simple method to simulate chemical reactions accurately is Gillespie’s stochastic simulation algorithm (SSA) [41], as used in some simulations [42].

#### Combinatorial complexity in models of CaMKII

One challenge in modelling CaMKII is that each CaMKII holoenzyme comprises multiple subunits; initial estimates were of 8-14 subunits, but EM and X-ray crystallography show that there are 12 subunits [43-45] arranged in two hexamer rings. Since a phosphorylated subunit can act as a kinase to its neighbour, the multiple subunits give rise to a combinatorially large number of meaningful configurations (states) of the holoenzyme. For example, a model with 6 subunits, each of which can be in one of 12 states, can be in 498,004 configurations according to the necklace function [46] and would therefore need the same number of ODEs to simulate. To simulate the dodecamer ring would require ∼ 10^12^ states, an impractical number of states to model with ODEs.

This combinatorial problem can be alleviated by model simplification, for example by (i) reducing the number of subunits to 4 and (ii) lumping together states that are invariant to rotations and adjusting the reaction rates between states according to their multiplicities [47]. These strategies are used in other deterministic and stochastic models of CaMKII [22, 48, 49]. A further simplification can be made by lumping together states with the same number of phosphorylated subunits, and weighting the transition rates between these states [39].

#### Agent-based simulation

Combinatorial complexity can also be dealt with using agent-based simulation, in which the states of individual molecules rather than populations of molecules are followed through the simulation [50]. For example, in simulations of a 10-subunit CaMKII holoenzyme [51], there was one variable per subunit, each of which described which of 5 states the subunit was in. The state of each holoenzyme was therefore described by 10 state variables, giving 976,887 states of the holoenzyme. Transition probabilities between a subunit’s states depended on its own state and that of its neighbouring subunit. Transitions were generated in 100 ms time steps in each subunit in turn, based on the state of the holoenzyme in the previous time step - similar to the *τ*-leap algorithm later formalised by Gillespie [52]. As this method is based on a fixed time step it can be combined with deterministic simulation of some elements of the system, as in a model of CaMKII activation in a dendritic spine [53].

#### Rule-based simulation

Agent-based simulation alone does not solve the problem of how to represent the states and the transitions between states clearly and concisely [50]. To specify transitions in agent-based simulations “rules” are specified in which the state of a fragment of system is mapped to the transitions that can occur within that fragment. For example a CaMKII monomer may be phosphorylated when both it and its neighbour (the fragment) are bound to Ca^2+^-CaM complex [24]. The StochSim agent-based simulator [54] describes rules by using flags to represent phosphorylation and binding states to be attached to molecules. However, the StochSim description of binding of CaM to CaMKII, phosphorylation states of CaM and trapping of CaM by CaMKII [24] is, arguably, unwieldy, requiring 1,209 lines of code.

Second generation rule-based modelling languages such as Kappa [55] or BioNetGen (BNGL, [56]) have a well-defined, general syntax to specify binding sites and states of proteins and interactions between protein binding domains. The interaction rules can be expanded to generate the “biological network”, i.e. the full set of complexes and reactions needed to simulate the system [56]. These reactions can be converted into ODEs or stochastic differential equations (SDEs), or simulated using a stochastic simulation method [41,52]. In another approach - dubbed “Network Free” [56], since no biological network is generated - simulators, such as KaSim [55] or NFSim [57], create the complexes that exist throughout a simulation dynamically. Network-free methods avoid the prohibitive memory requirements needed to store all possible states in a large network [57], and even allow simulations with infinite numbers of potential species [55]. This form of “on-the-fly” simulation is intrinsically stochastic, with transitions occurring one rule at time, similar to Gillespie’s SSA [41]. For smaller networks, ODEs, SDEs or the SSA are faster, but because the simulation speed of these methods scales roughly with network size (i.e. the number of reactions), for larger networks these conventional methods are slower than network-free simulation [57].

#### Varying model structures

Authors devise differing descriptions of the same pathway. For example Byrne et al. [58], Stefan et al. [59] and Faas et al. [14] all describe the binding of calcium to CaM, but each model has a distinct structure. The models of Byrne et al. and Faas et al. assume cooperativity within the N and C lobes of CaM: the rate at which a calcium ion binds to a lobe with one calcium bound is different from the rate at which calcium binds to the lobe in the *apo*, unliganded, state. In contrast, Stefan et al. assume that the affinity of each of the four positions on CaM is independent, but that these affinities depend on whether the entire CaM molecule in the “tense” or “relaxed” conformation [60], which is an allosteric mechanism [61]. The two positions within each lobe are assumed to be equivalent by Faas et al., but not by Byrne et al. The model of Faas et al. has been fit against kinetic data, which is richer than the binding curves fit by Byrne et al. and Stefan et al., but it has not been investigated whether the parameters of these earlier models could be adjusted to fit the kinetic data.

There is also diversity in the number of states monomers in models of CaMKII may assume, and how the multimeric structure of the molecule is represented. An additional variation in particle-based simulations of CaMKII is that once the CaM N or C lobe is bound to a CaMKII monomer, it becomes much more likely that the other lobe on the same CaM molecule will bind to a neighbouring CaMKII monomer on the hexamer ring [58]. This necessary to fit Ca-chelator-induced dissociation curves [62] and steady-state CaM-CaMKII binding curves [43]. A result of this assumption is that the rate of CaM binding to CaMKII is dominated by the more affine N-lobe.

#### Biophysical constraints on parameters

A number of strategies are used to reduce the considerable number of reaction coefficients in molecular models. For example, the reactions in Byrne et al. [58] are parameterised by 2 sets of 24 parameters, but the forward reaction coefficients are all set to be equal, reducing the number to 2 sets of 13. The principle of microscopic reversibility [61] is used to link reaction coefficients that are in loops, taking the number down to 2 sets of 9. Microscopic reversibility applies generally, though some ion channels are exceptions to this rule [61]. Other linkages between parameters can be postulated; for example in the allosteric model of Stefan et al. [59], the ratio between the affinities of each site for calcium in the tense and relaxed conformations is assumed to be the same for each of the four sites.

#### Data used to constrain parameters

Various types of data have been used to constrain the parameters of single pathway models. To obtain equilibrium binding curves, equilibrium dialysis with radioactively labelled ligands can be used, as by Crouch and Klee in their determination of Ca^2+^-CaM binding. More recently, stopped-flow fluometry [43] has been used for the same purpose. This method has the disadvantage of a relatively long dead time of the order of 2 ms, which hinders determining fast dynamics, e.g. of the N lobe of CaM. A faster method is calcium uncaging, which can lead to a sub-0.1 ms change in calcium concentration, and measurement with a fast fluorescent calcium indicator [14].

Spectroscopic analysis can be used to infer conformational changes, e.g. the tense to relaxed conformation change upon binding of a calcium ion to CaM [60]. Phosphorylation states, e.g. of CaMKII, can be measured using radioactively labelled ATP [43] which can be coupled with immunoprecipitation and gel electrophoresis [63].

#### Optimisation of free parameters

Even after reducing the number of parameters there are typically a number of free parameters in a model, and a number of optimisation techniques are used to fit them to data, for example particle swarm optimisation [58]. Latin hypercube sampling can be used to determine global parameter sensitivity [20].

#### Hypothesis-driven and simplified modelling

In one combined experimental-modelling study [63], the authors engineered a monomeric form of CaMKII. This allowed them to measure the CaM-dependent phosphorylation properties of CaMKII and produce a simplified computational model, which predicted that the amount of CaMKII activation would depend on the frequency of a presented train of Ca pulses: CaMKII could thus act as a frequency decoder. A number of CaMKII models at various levels of detail have been formulated to explain the dependence of CaMKII activation on the frequency of calcium pulses [47, 48, 64].

#### Data-driven rule-based modelling

Proteomic studies of the synapse (Table S2) show that there are many proteins in the synapse not included in the models described thus far. The challenges of combinatorial complexity, already encountered in models of CaMKII, are magnified as more proteins are added. Rule-based modelling has been applied to simulate a network containing 54 proteins, with interactions were described by 136 rules [23]. This model makes predictions about the molecular composition of complexes that could occur in the PSD.

### Spatial models

The modelling methods described so far assume that molecules are within a well-stirred, spatially homogeneous environment. However, the cellular environment is not homogeneous; for example, calcium enters through N-methyl-O-aspartic acid receptors (NMDARs) on one side of the spine head. It can react with buffers on a shorter timescale than it takes to diffuse through the spine, and can exist within microdomains around the NMDARs briefly at high concentrations. Thus, to address some questions, it is necessary to model space explicitly.

#### Deterministic reaction-diffusion

Deterministic diffusion is modelled by splitting cellular space into compartments and formulating ODEs to describe how reactions within compartments and fluxes between compartments affect the concentrations of species within each compartment. Deterministic diffusion along one dimension has been used in models of calcium and other intracellular signalling in spines [65-67]. Whilst these models do not model LTP and LTD explicitly, they give insights such as that the combination of calcium pumps and buffers can confine calcium and activated CaMKII to the synaptic spine head [67], or that the temporal ordering of input at weak and strong synapses with NMDARs determines the concentration of calcium in the spine, which will then influence the intracellular pathways underlying LTP and LTD [66]. The NEURON simulator, used widely in models of electrical activity of neurons, also supports reaction-diffusion, with recent work to extend these capabilities [68]. Deterministic reaction-diffusion can be simulated in 3D by splitting cellular space into tetrahedral or cubic compartments, as implemented in the STEPS simulator [69].

#### Compartmental stochastic reaction-diffusion

The numbers of molecules in each compartment of a mesh is often small enough to warrant stochastic simulation methods. Gillespie’s SSA can be extended to a compartmentalised volume by replicating the set reactants in each compartment, and treating diffusion of reactants between each compartment as a type of reaction [41]. This “Spatial SSA” method and more efficient approximations [70] have been used for a number of simulations of medium spiny projection neurons in the striatum [28-31] and is implemented in the simulators NeuroRD [28] and STEPS [69].

#### Compartmental agent-based stochastic reaction-diffusion

The Spatial SSA requires one variable in each compartment to describe the number of molecules in every possible state in the system, and therefore is ill-adapted to deal with models of molecules with many states, such as CaMKII. A custom extension to the Spatial SSA has been used to study the relative effects of the stochastic opening and closing of NMDARs and of stochastic binding between CaMKII holoenzymes and CaM in a spine head [27]. The results showed that NMDARs were a greater source of noise, due to their smaller numbers than the CaMKII holoenzymes. The agent-based, rule-based simulator SpatialKappa [71] extends the Kappa language syntax and the KaSim algorithm to allow diffusion of complexes between voxels in regular meshes.

#### Particle-based stochastic reaction-diffusion

In particle-based simulation methods, each molecule has a location in 3D space or on a 2D membrane and moves in Brownian leaps. Reactions may occur when particles come within an interaction radius of each other. Simulators implementing this method include MCell [72] and Smoldyn [73]. MCell has been used to model diffusion of glutamate molecules in the synaptic cleft and their binding to NMDARs and *α*-amino-3-hydroxy-5-methyl-4-isoxalone propionic acid receptors (AMPARs) [27,74], and influx of calcium into the spine head and its interaction with calcium binding proteins [75, 76]. The most recent version of Smoldyn supports the rule-based BNGL language, but only to generate reaction networks, not to perform network-free simulation.

#### Modelling diffusion measurements

Khan et al. [32] used a spatial model built with Smoldyn to interpret their fluorescence recovery after photo-bleach (FRAP) measurements of CaMKII diffusing in a spine head before and after glutamatergic stimulation. Eleven bidirectional reactions described binding of phosphorylated CaMKII to the PSD, binding of non-phosphorylated CaMKII to the actin cytoskeleton, and CaMKII self-aggregation. All these reactions contribute to keeping stable CaMKII concentrations in stimulated spines, providing an explanation of sequestration of CaMKII in dendritic spines.

#### Multiscale modelling

It is possible to simulate reaction-diffusion and the membrane potential using the same spatial mesh, but these simulations are likely to run very slowly because of the unnecessarily fine mesh in parts of the model, such as the dendrites, where concentration gradients are lower. Multiscale modelling, defined as the process of using multiple models at different scales simultaneously to describe a system [77], can allow for the desired level of detail with tractable simulation times. To demonstrate a multiscale algorithm to integrate detailed models of signalling networks within electrical models of neuron, Mattioni and Le Novère [34] used a model of a striatal medium spiny projection neuron (MSPN) with 1,000 synaptic spines attached. The electrical activity and calcium accumulation in the dendrites and soma of the neuron were simulated using the NEURON implementation of the compartmental modelling method. Within each spine, the calcium flux through AMPARs, NMDARs and voltage gated calcium channels (VGCCs) calculated by the electrical model is fed to instances of a molecular simulator (in this case E-CELL3), in which the calcium binds to CaM, which then participates in a biochemical network typical of striatal MSPNs. A similar effort has incorporated the rule-based SpatialKappa simulator into NEURON [78].

### Models of hippocampal synaptic signalling pathways

In tandem with the extensive experimental study of LTP and LTD in the hippocampus, computational models of hippocampal synaptic plasticity have been developed.

#### The CaMKII phosphorylation-dephosphorylation circuit

Lisman [79] proposed a model to account for how LTP and LTD could be mediated by postsynaptic calcium acting as a second messenger (Fig 1). A high concentration of calcium, caused by coincident pre-and postsynaptic activity, leads, via binding to CaM, to phosphorylation and then auto-phosphorylation of CaMKII. At moderate concentrations calcium binds to calcineurin (PP3), which is also known as PP2B; we use PP3 for consistency with gene identifiers. The calcineurin-calcium complex dephosphorylates protein phosphatase inhibitor 1 (I1), thereby deactivating it. The inactive I1 then unbinds from protein phosphatase 1 (PP1), allowing it to dephosphorylate phosphorylated CaMKII. At high Ca^2+^ levels this pathway is inhibited via Ca^2+^-CaM activated adenylate cyclase (AC), which then catalyses production of cyclic adenosine monophosphate (cAMP) from adenosine triphosphate (ATP). The cAMP then binds to the regulatory subunits of cAMP-dependent protein kinase (PKA), releasing its catalytic subunits which then phosphorylate I1, thereby allowing it to sequester PP1. The Ca^2+^-CaM complex also activates phosphodiesterase (PDE), which hydrolises cAMP into adenosine monophosphate (AMP), thus reducing the rate of activation of PK A.

**Fig 1.**
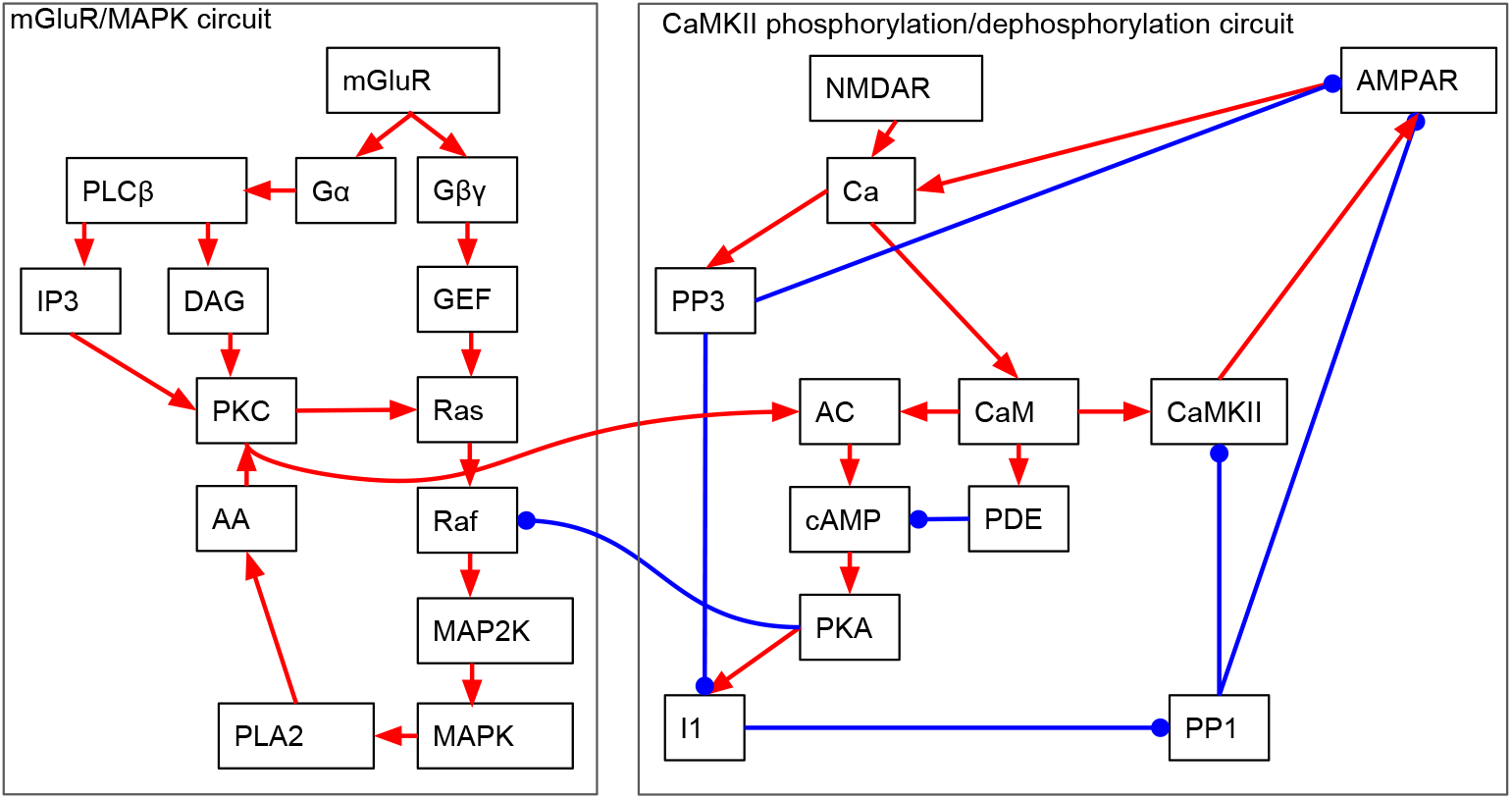
Partial block diagrams comparing essential elements of the hippocampal biochemical circuit. Each small box represents an ion, monomer or multimer. Red arrows indicate activating interactions. Blue lines ending in circles represent inhibiting interactions. Within each box, the molecules can be one of potentially many binding or phosphorylation states. The circuit is split into two sub-circuits: the CaMKff phosphorylation/dephosphorylation circuit and the mGluR/MAPK circuit.

Lisman formulated this biochemical circuit as a simplified steady-state mathematical model of the net phosphorylation rate of CaMKII, and showed that a set of parameters existed that would allow unphosphorylated (“off”) CaMKff molecules to be phosphorylated (activated) by high Ca^2+^ levels, and phosphorylated (“on”) CaMKff molecules to be dephosphorylated (inactivated) by low Ca^2+^ levels. Lisman hypothesised that, ultimately, CaMKII activation increases the non-NMDA component of the synaptic response. The biochemical circuit of Lisman is included in a number of dynamical biochemical models of postsynaptic signal transduction [80-84]. In some cases PKA is assumed to be tonically active rather than released from inhibition by cAMP, [83, 85] and other features may be included such as sequestering of CaM by neurogranin and SAP97 [83].

#### AMPA receptor phosphorylation

Models have been formulated in response to the developing understanding of AMPARs [86]. AMPARs comprise four subunits, each of which is one of GluR1-4. The phosphorylation at two sites on GluR1 affects the function of the AMPAR multimer. In synapses in a “naive” state, i.e. those which have not been exposed to any plasticity protocols, phosphorylation of Serine 831 (Ser831), by CaMKII or protein kinase C (PKC), is associated with LTP [87, 88] and dephosphorylation of Serine 845 (Ser845) is associated with LTD [89]. In synapses that have already experienced LTD, “dedepression” caused by a theta-burst stimulus is associated with Ser845 phosphorylation, and in a synapse that has potentiated, the Ser831 site is dephosphorylated during “depotentiation” [88].

These findings led to the four state model of AMPARs by Castellani et al. [90], in which potentiation is caused by phosphorylation of the Ser831 and Ser845 sites, and LTD caused by dephosphorylation of the sites. The activation of the phosphatases and kinases was set up in the model so that the phosphates were more activated than the kinases at low concentrations, and vice-versa for high concentrations. Steady-state analysis of the set of 4 bidirectional reactions gave a typical biphasic Ca^2+^-synaptic strength curve in which there is LTD at moderate concentrations of calcium and LTP at high concentrations. Furthermore, control of Ca^2+^ levels via adaptation of NMDARs allowed modification of the threshold level of Ca^2+^ at which LTP rather than LTD occurred, as in the Bienenstock-Cooper-Munro (BCM) rule [91].

#### AMPAR trafficking

Blocking AMPAR exocytosis causes run-down of synaptic strengths, and inhibiting endocytosis of AMPARs causes an increase in AMPAR responses [92]. This discovery lead to the idea of a stable distribution of receptors at the synapse being replaced by a highly dynamic picture, with continuous exocytosis and endocytosis of AMPARs [93]. The trafficking to synapses comprises three steps [94]: (1) AMPARs bound to Transmembrane AMPA receptor regulatory protein (TARP) proteins such as stargazin are inserted into the dendritic shaft or spine by phosphorylation events caused by PKA, PKC, extracelluar regulated kinase (ERK) (part of the mitogen-activated protein kinase (MAPK) family) or Phosphoinositide 3-kinase (PI3K), or myosin-V; (2) the AMPARs diffuse through the membrane to the synapse; and (3) phosphorylation events (triggered by active CaMKII targeting stargazin) increase the affinity of the AMPAR-stargazin complex for PDZ-containing scaffolding proteins such as PSD95, PSD93, SAP97 and SAP102. AMPAR trafficking away from synapses is thought to be an inverse process, whereby AMPARs are released from PDZ proteins and diffuse from the synapse back to the dendrite, where they are endocytosed. There is a link between trafficking and the phosphorylation states of AMPARs, with phosphorylation of Ser845 on the GluR1 subunit needed to incorporate GluR1 subunits into synapses [95], although it is not clear how strong this link is [96].

#### Integrated modelling of CaMKII phosphorylation circuit and AMPAR trafficking

Urakubo et al. [84] explored whether a model that integrated AMPAR trafficking with the CaMKII-phosphorylation-dephosphorylation biochemical circuit first formulated by Lisman and implemented by Bhalla and Iyengar [80] could account for spike-timing dependent plasticity (STDP). They embedded the circuit in a spine containing NMDARs, AMPARs and VGCCs in a simplified soma-and-dendrite compartmental model with conductances used in models of CA1 hippocampal cells [97, 98]. In their first model LTP resulted from pre-before-post spiking, but LTD did not result from post-before-pre spiking. To cause LTD in this situation, it was sufficient that the NMDARs were blocked by binding of Ca^2^+-bound CaM. This biochemical detection circuit was linked to AMPAR phosphorylation and dephosphorylation by the activities of the kinases CaMKII and PKA and the phosphatases PP1, PP3 and protein phosphatase 2 (PP2) (commonly known as PP2A). AMPAR trafficking was modelled by having four pools of AMPARs: (1) cytosolic; (2) in the dendritic or spine shaft membrane; (3) at the synapse but not anchored by PDZ proteins; and (4) at the synapse, anchored by PDZ proteins. The phosphorylated LTP and LTD states were used to control the rates of endo-and exocytosis, and binding to the PDZ proteins.

#### The MAPK circuit and metabotropic glutamate receptor (mGluR) signalling

To the CaMKII phosphorylation-dephosphorylation circuit the modular model of Bhalla and Iyengar [80] adds the MAPK cascade, activated by mGluRs (Fig 1). Input to mGluRs activates G-proteins, which then go on to activate phospholipase C-ß (PLC-ß), leading to production of diacylglycerol (DAG) and inositol (IP3) and phosphorylation of PKC. This activates the cascade of Ras, Raf, mitogen-activated protein kinase kinase (MAP2K) and MAPK. In turn, MAPK activates phospholipase A2 (PLA2), which cleaves arachidonic acid (AA) from phospholipids. The AA binds to PKC, activating it, which in turn leads to more Ras activity, completing the loop. The G-proteins also activate the Ras-Raf-MAP2K-MAPK pathway via up-regulation of guanine exchange factor (GEF). The parameters in the system were such that the persistent up-regulation of PKC was enough to catalyse AC production in the CaMKII circuit, and thus up-regulate PKA and down-regulate PP1, leading to prolonged CaMKII activation. There was also inhibitory crosstalk from the CaMKII to the MAPK via inhibition of Raf by PKA.

#### Late LTP, synaptic tagging and gene expression

The models described so far all deal with the induction of early-LTP, which occurs up to 4 hours after induction and does not depend on protein synthesis [99]. In contrast, late-LTP depends on protein and mRNA synthesis. In order to solve the conundrum of how AMPAR proteins, which were assumed not to be synthesised close to synapses, get to the synapses, Frey and Morris [99] proposed that a “synaptic tag” is set when activity has potentiated the synapse. Smolen et al. [100] formalised this concept into an ODE model containing four pathways: (1) the MAPK cascade; (2) PKA activated by cAMP; (3) CaMKII; and (4) Ca^2+^-activated calcium/calmodulin-dependent protein kinase kinase (CaMKK), which activates calcium/calmodulin-dependent protein kinase (CaMKIV). The CaMKII, MAPK, and PKA pathways are all required to set a synaptic tag. CaMKIV, assumed to be in the nucleus, and MAPK are assumed to activate unknown transcription factors. The input to the model was the assumed time courses of Ca^2+^, Raf and cAMP. The CaMKII phosphorylation circuit was not modelled.

To induce late-LTP, translation and synaptic tags need to be active simultaneously. Smolen et al. [16] devised a distinct model at a similar, relatively low, level of detail containing notional synaptic LTP tags activated by Ca^2+^-CaM-CaMKII, LTD tags activated by the Raf-MAPK pathway, local protein translation mediated by autonomously active isoform of atypical protein kinase C ζ (PKMζ) (after a chequered history, back in favour as a memory molecule [101]), and movement of PKMζ and notional plasticity related proteins from the cytoplasm to synapses. The model was used to explore how strong potentiating or depressing stimuli at one synapse can promote protein synthesis that allows, at other synapses, weak stimuli to cause plasticity.

### Models of striatal synaptic signalling pathways

The striatum integrates multiple inputs to the basal ganglia, such as glutamatergic excitatory afferents from the cortex and dopaminergic inputs from the midbrain [102]. Around 95% of striatal cells are MSPNs, in which signalling cascades activated simultaneously by glutamatergic and dopaminergic stimuli is a necessary condition for the LTP that underlies reinforcement learning [103]. Models of striatal MSPNs share some pathways with hippocampal synapses and include striatum-specific proteins.

#### Multistate DARPP-32

An abundantly expressed protein in MSPNs is phosphatase 1 regulatory subunit 1B (PPP1R1B), known as dopamine-and cAMP-regulated neuronal phosphoprotein with molecular weight 32 kDa (DARPP-32). As a homologue of I1, it has the same major role of PP1 inhibition. It is a hub protein that is regulated by multiple neurotransmitters and phosphorylation sites. There are at least 8 modification sites known in the DARPP-32 amino acid sequence, and 4 of them are known to have a regulatory impact on DARPP-32 [104]. The threonine sites (Thr34 and Thr75, as positioned on the rat protein sequence) have a major regulatory role in signal processing. Thr34 inhibits PP1 and is phosphorylated by PKA, which Thr75 inhibits. The serine sites (Ser137, Ser102, as positioned on the rat protein sequence) regulate Thr34 positively. Ser137 inhibits dephosphorylation of Thr34 on Ca^2+^ stimulation and Ser102 enhances phosphorylation of Thr34. A number of models of dopamine (DA) and Ca^2+^ signal integration have included only Thr34 and Thr75 as major switching factors between LTP and LTD [17,18,29,105]. A few models incorporate all four phosphorylation sites [19,106].

#### Glutamatergic and dopaminergic signal integration

Lindskog et al. [105] created an ODE model of interacting cascades activated by DA and Glutamate (Glu) signals stimulating dopamine receptor D1 (DRD1) and Ca^2+^ influx through NMDAR, respectively. The glutamatergic signalling cascade shares the general network structure of the CaMKII circuit with hippocampal models (Fig 2), with a few major differences.

**Fig 2.**
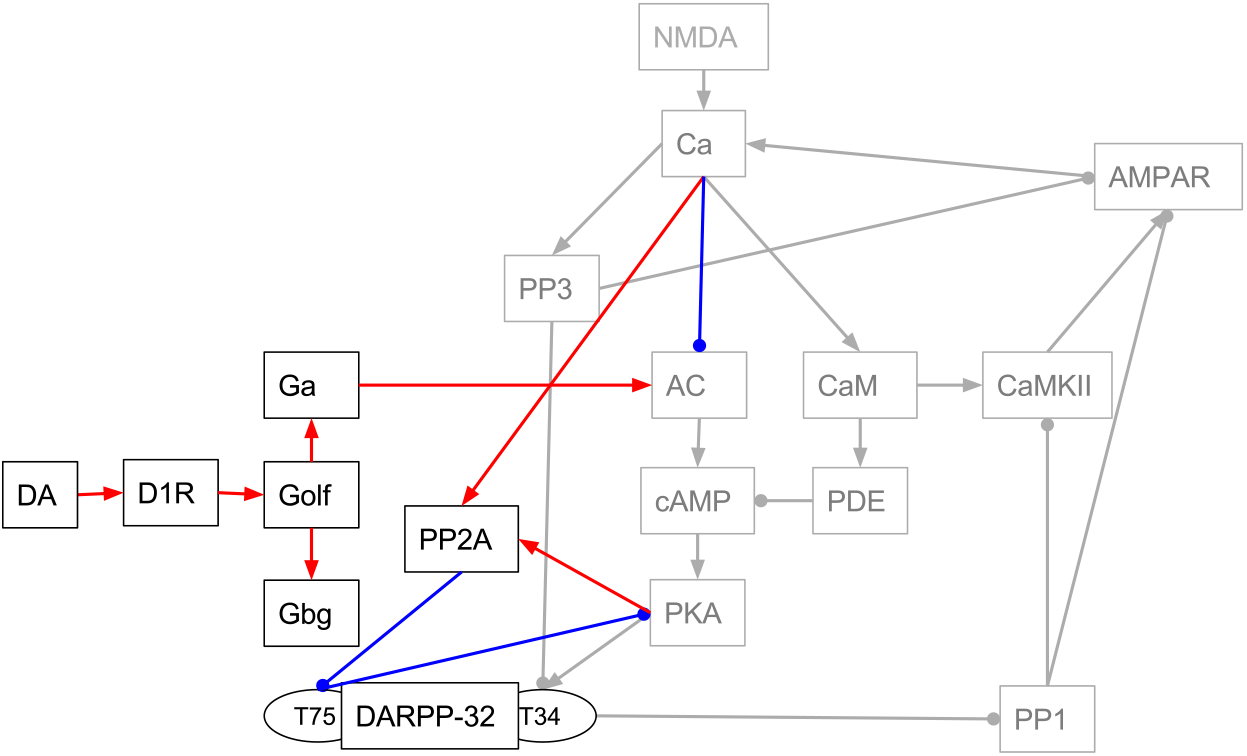
Incomplete block diagrams comparing essential elements of striatal biochemical circuit. Greyed nodes and edges denote shared elements with hippocampal models. See Fig 1 for explanation.

Firstly, the inhibition of PP1 does not occur via I1 but rather via DARPP-32 phosphorylated at Thr34. Secondly, as the DRD1 is a G-protein-coupled receptor (GPCR), DA input adds to the network G-protein activation events. On DA stimulation, G*_αβγ_* dissociates into G_*α*,olf_ and G_*βγ*_ subunits. Subsequently, G_*α*,olf_ binds to AC and ATP, synthesising cAMP. The last event, which results in activation of PKA and the cascade inhibiting PP1, is shared by both hippocampal and striatal models. However, in contrast to hippocampal models, in Lindskog’s model [105], Ca^2+^ inhibits AC, leaving its activation to DA input. Furthermore, Ca^2+^-activated PP3 dephosphorylates Thr34 counteracting the DA, but not the Ca^2+^ signal.

In the model Thr34 is both activated and inhibited by a Ca^2+^ feedforward signal, which is conveyed by the PKA-PP2-Thr75 double negative feedback loop. PP2 dephosphorylates Thr75 but its action is enhanced by Ca^2+^ and PKA. The model showed that the loop does not exclusively reinforce PKA pathway stimulated by DA but instead acts as a competitive inhibitor for PKA.

The detailed model of Nakano et al. [15] demonstrated that the loop can have a major role in LTP induction. They extended the network upstream of DARPP-32 and added AMPAR phosphorylation and trafficking as a direct readout of plasticity. Their model required activation of both CaMKII and PKA to reach striatal LTP. They also included the downstream pathway of mGluR activation that represented mainly the bi-directional effect of Ca^2+^ on IP_3_ receptor located at the endoplasmic reticulum.

#### STEP-mediated crosstalk between glutamatergic and dopaminergic signalling cascades

Gutierrez-Arenas et al. [18] developed a signalling model of two main signalling pathways activated by DA and Glu inputs in MSPNs: AC-cAMP-PKA and NMDAR-Ca^2+^-Ras. The AC-pathway was built on the model of Lindskog [105] by adding a NMDAR-cascade, in which the dissociated G_*βγ*_ subunits activate Fyn which phosphorylates a NMDAR subunit, thus enhancing the Ca^2+^ influx. Ca^2+^ activates the MAPK pathway phosphorylating mitogen-activated protein kinase 1, also known as ERK2 (MAPK1) at two sites. In striatal plasticity, MAPK1 activation is known to require both DRD1 and NMDAR stimulation, as shown by the negative impact on MAPK1 phosphorylation in the DARPP-32-knockout mouse model [107]. DRD1 activation by the DA-signal also enhanced the Ca^2+^ current through NMDAR, e.g. by the phosphorylation of NMDAR by activated PKA. This particular reaction network was chosen to allow for examination of various scenarios that could explain the results of behavioural experiments showing distinctive segregation of behaviours of two animal types representing G_*α*,olf_-deficiency and DRD1-deficiency. The former exhibited disruption of phosphorylation of the GluR1 subunit of AMPAR and the latter disrupted phosphorylation of MAPK1 after acute psychostimulant administration. This effect was present despite known crosstalks between two cascades mediated by striatal enriched tyrosine phosphatase (STEP), which could balance the sensitivity in both pathways. The model reproduced the segregation with an assumption that there are two DRD1/G_olf_ signalling compartments for each pathway distributed from common pools of DRD1 and G_olf_. These compartments differ in DRD1 and G_olf_ distribution determined by the opposite affinity strengths for these molecules in each compartment. These settings resulted in a competition between the two compartments for G_olf_ /DRD1 resources, giving a ‘winning hand’ to the one with a stronger affinity to a given molecule.

#### Interactions between G-protein-coupled receptors

DRD1 is a subfamily of dopamine receptors and one of multiple types of GPCRs expressed in MSPNs, including serotonin (5-HT_2*C*_ receptor [108]), noradrenaline (*α*_2_-adrenoceptor, *β*_1_-adrenoceptor [109], acetylcholine (muscarinic M4 receptor; M4R), adenosine (A2a receptors; A2aR) and dopamine receptors of D_2_-like family. The last three, alongside DRD1, were modelled by Nair et al. [17], who simulated the reward prediction error (defined as the difference between the received and expected reward). They modelled two types of MSPNs, expressing either DRD1 and M4R (striatonigral projections) or DRD2 and A2aR (striatopallidal projections). These two types of neurons process DA-signals in two opposing manners by stimulating (DRD1-expressing) or inhibiting (DRD2-expressing) the signalling cascade resulting in phosphorylation of DARPP-32 at Thr34. In both models neuromodulators interact through G_*i*/*o*_ and G_olf_ signalling, inhibiting and activating AC5 respectively. Also in both models, AC5 is inhibited by G_*i*/*o*_ at the basal state. In the DRD1-expressing neurons, G_*i*/*o*_ is coupled with the M4R-tonic ACh signal; and in the DRD2-type of neurons with the DRD2-tonic DA signal. In DRD1-neurons, the high PKA activation level was achieved with a simultaneous DA-peak and ACh-dip. These neurotransmitter signals realise an AND-gate, sensitive but noise-prone to a positive reward. In DRD2-neurons it is the DA-dip that increases the PKA activation, even without Adn signal. This suggests that in this type of neurons the cAMP-PKA cascade mainly detects reward omission.

#### Spatial specificity in synaptic plasticity

The model of Oliveira et al. [29] studied the mechanisms of spatial restriction of PKA activation by A-kinase anchoring protein (AKAP). The problem required a multi-compartmental stochastic reaction-diffusion approach. To evaluate distinct functions of anchoring, the experimental protocol consisted of four spatial variations in localisation of AC and PKA, either locating them in the spine head or at dendritic submembrane area. The signalling network was adopted from Lindskog [105] and the stimulating signal was either dopamine alone, corresponding to the reward response, or the combined DA and Ca^2+^ influx used for LTP protocols. The results showed that for the induction of LTP the colocalisation of PKA near the source of cAMP is more important than its colocalisation near its target substrates (e.g. DARPP-32, PP2, PDE).

Kim et al. [31] used the NeuroRD algorithm to model 19 molecules in the postsynaptic signalling pathways of the dendrites of striatal MSPNs with multiple spines. The model investigated the hypothesis that temporal patterns, linked to Ca^2+^, determine LTP or LTD induction, via PKC or endocannabinoid 2-arachidonoyl-glycerol (2AG) production respectively. The ratio between the number of activated PKC and 2AG molecules was used as an indicator of the direction of plasticity. It describes G_q_-coupled pathways, the temporal pattern of Ca^2+^ stimulation and G_a q_ activation. In the simulations LTP was specific to spines, whereas LTD was more diffuse. This suggested that spatiotemporal control of striatal information processing uses Gq-coupled pathways for decision-making.

### Cerebellar synaptic models

Despite the historical importance of cerebellar granule cell to Purkinje cell plasticity, at least 9 types of synaptic and non-synaptic plasticity are known [110]. The classical LTD at cerebellar granule cell to Purkinje cell synapses occurs when there is simultaneous climbing fibre and granule cell (parallel fibre) firing. At the heart of the model of Kuroda et al. [111] is the MAPK positive feedback loop found in hippocampal and striatal models [18,80], which here comprises Raf-MAP2K-MAPK-PLA2-AA-PKC. Parallel fibre activity both activates and inhibits the loop. Parallel fibre glutamatergic input to AMPARs causes Na+ influx, which triggers the Na^+^/Ca^2+^ exchanger causing Ca^2+^ influx which, in turn, activates PKC and PLA2. PKC is also activated via mGluR and AMPARs also activates Lyn tyrosine kinase directly, which activates Raf in the MAPK loop. Parallel fibre input also releases NO, which, via the guanylate cyclase-cGMP-PKG pathway, activates PP2, which inhibits MAP2K. Climbing fibre inputs also activate the MAPK link via Ca +, and via Raf which is activated corticotropin releasing hormone receptors (CRHR) activated by corticotropin releasing factor. When the loop is active, activated PKC phosphorylates AMPARs, but in contrast to hippocampal models phosphorylated AMPARs are internalised, leading to LTD.

Antunes and DeSchutter [112] model LTD in cerebellar granule cell to Purkinje cell synapses in the cerebellum using Gillespie’s SSA, as implemented in the STEPS simulator. The model includes a version of the PKC-MAPK circuit (Fig 2), but with an undetermined “Raf-activator” between PKC and Raf. This Raf-activator could be Ras itself or indirect activation of Ras via complex Src/Proline-Rich Tyrosine Kinase 2 (PYK2). PP5 tonically inhibits Raf and MKP (DUSP) inhibits MAPK. Activated PKC promotes endocytosis of AMPARs, thus causing LTD. The stochastic nature of the model leads to LTD being stochastic and binary at individual synapses, but over the ensemble of synapses this results in a graded relationship with the magnitude of the activating Ca^2+^ signal. Increasing the number of molecules makes the system less stochastic, and makes the resulting macroscopic signal less graded.

Antunes et al. [42] extend this model by incorporating CaMKII and PP3 to implement LTP. They use the rule-based BioNetGen system to generate stochastic reactions that are simulated using Gillespie’s SSA. In contrast to hippocampal models, calcineurin promotes LTP by preventing endocytosis of AMPARs. RKIP is also incorporated as an additional activator of Raf.

### Summary

In summary, the development of biophysical models of synaptic plasticity has been propelled by: (1) hypothesis-driven physiological and molecular biological discoveries; (2) the need to formalise informally expressed hypotheses; (3) the intrinsic fascination and intellectual challenge of complex biomolecules such as CaMKII; and (4) increasing compute power, which makes it practical to model stochastic and spatial aspects of synaptic signalling cascades. Challenges in the field have included dealing with combinatorial complexity and finding appropriate sets of parameters. Recent computational modelling methods, such as agent-based and particle based simulation, address the problem of computational complexity. Depspite being an active field of research, the perennial problem of inferring parameter values remains more intractable.

## Analysis of proteins in synaptic models

Computational models of synaptic plasticity are important tools for understanding synaptic and neural function. When they include molecular entities and phenomena they can also be used to study dysfunction, and potentially model pharmacological interventions. Clearly the coverage of synaptic molecules found in the existing ‘model space’ is going to be very incomplete given the intense amount of effort required to develop each model but here we sought to explore systematically molecular coverage to identify significant gaps that might offer new opportunities.

Computational models contain a diverse cast of players, including proteins, second messengers, reporters, ions and others. Models vary in how precisely they specify proteins; for example Bhalla and Iyengar [80] specify AC1, AC2 and AC8, whereas Castellani et al. [82] and Oliveira et al. [28] specify AC, which could, in principle, map to any of the adenylate cyclases expressed in the synapse. This presents a problem when mapping models to molecular identifiers, which we addressed by developing a mapping from what we refer to as model “entities” to gene families. For example a protein such as Calmodulin 1 can be mapped onto a single gene (*CALM1*), but a family of proteins such as metabotropic glutamate receptors maps onto more than one gene (*GRM1*-*GRM8*). By definition, second messengers or ions do not map onto gene symbols.

The concept of entities allows each model’s constituents to be catalogued faithfully and then mapped onto identifiers according to the steps shown in Fig 3: (1) select models to analyse; (2) determine all entities (e.g. proteins, protein multimers or families, ions and second messengers) that are contained in each model; (3) map these entities onto gene identifiers and higher level families; and (4) use the lists of entities in each model and the mappings to undertake comparative analyses. These analyses include: comparison of modelled proteins with pre-and postsynaptic proteomic dataseis; identification of properties of modelled genes, in particular cellular pathways, gene ontology terms and disease; and comparison of models with each other.

**Fig 3.**
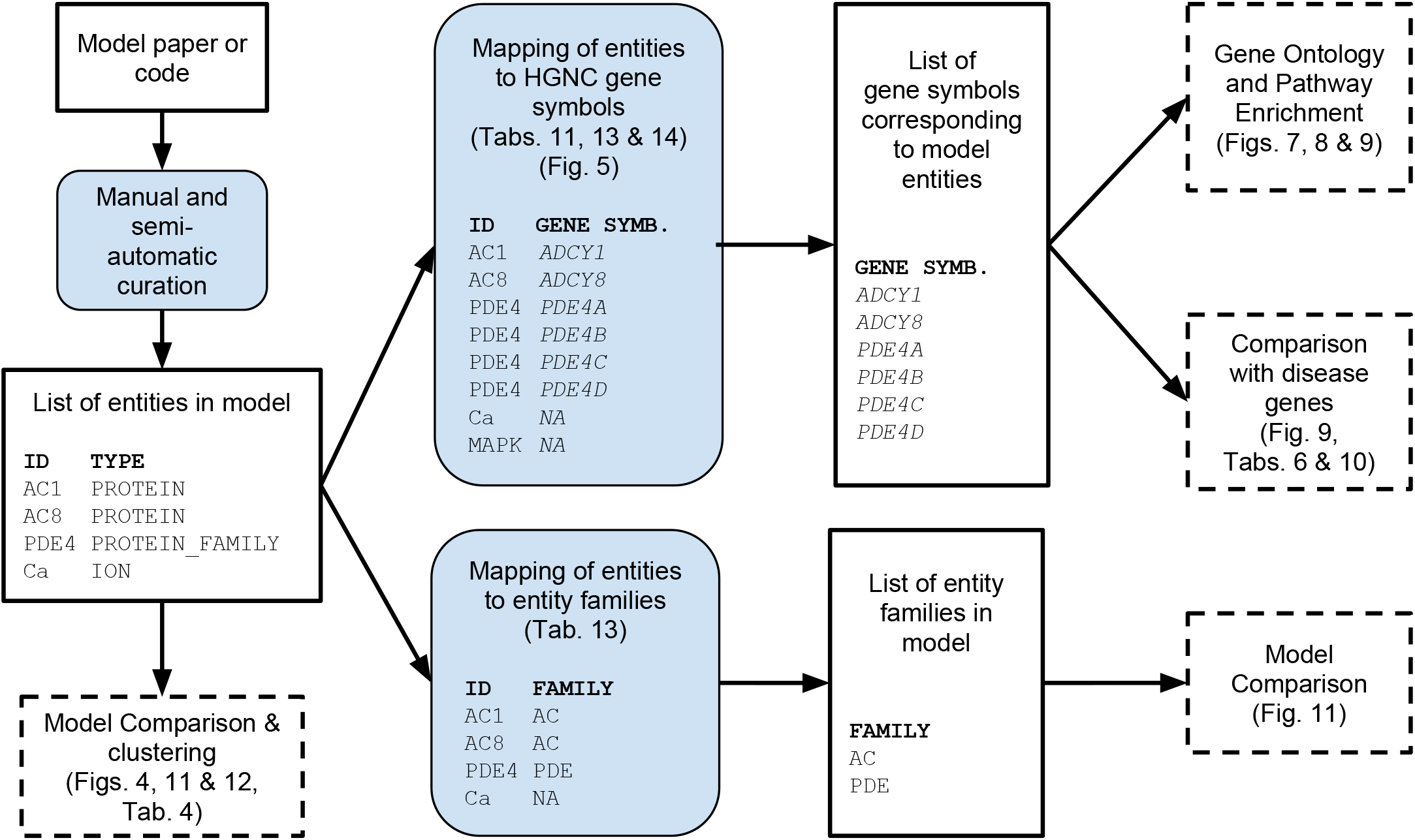
Overview of the modelling paper analysis process. Sets of data are shown in boxes with black rectangular in boxes with blue backgrounds and curved corners. Final analyses are shown in boxes with i the modelled entity. Boldface type refers to column headers.

### Selection of models

We selected a number of published computational, biophysical models of synaptic plasticity or related pathways (Table 3). Models that we regarded as phenomenological or descriptive, i.e. models describing a function with no explicit reference to an underlying mechanism, were excluded. For example, models of spike-timing dependent synaptic plasticity are phenomenological, since they contain an empirical function that maps spike times onto changes in plasticity with no reference to proteins.

The process of identifying the model constituents can be time-consuming, especially when machine-readable descriptions are not available. In order to address our questions regarding the molecular coverage of synaptic models, it sufficed to select a set of models that we were reasonably confident gave good genetic coverage, rather than to identify entities in every model. We assessed molecular coverage of pre-2010 models from the tables in Manninen et al. [9] and we screened models published between 2010 and December 31st 2015.

### Sources of models

A number of the models we selected are written in standardised modelling languages and hosted in large scale repositories such as ModelDB [113], BioModels [114], DOQCS [115] and the CellML repository [116]. ModelDB is a curated database of computational neuroscience models at the molecular and electrophysiological levels, written in a number of languages. BioModels hosts models which focus on biochemical and cellular systems at the physiological and biochemical levels, unrestricted by the biological subject [114,117]. In the curated branch of BioModels, models have to be annotated according to the minimal information requested in the annotation of biochemical models (MIRIAM) standard [118], thus meaning that model constituents are mapped to external identifiers. CellML is both a model format and a repository. The repository hosts a wide range of biological models, which have documentation pages generated from the meta-data supplied by model authors. DOQCS (Database of Quantitative Cell Signalling) is a database tailored for storing chemical kinetics and reaction level information [115]. The chemical-level description of each model corresponds to the GENESIS/Kinetikit simulator and reflects reaction diagrams or ODE equations.

Table 2 summarises the numbers of models we analysed that are stored in repositories and other locations, and the format of the model descriptions. Three of the 7 models deposited in the BioModels database were curated to MIRIAM standards. Around half of all catalogued models (14) had non-machine readable descriptions. Models in this group are often difficult to explore and extract information proves challenging. There were 18 machine-readable models available from publication attachments, on institute or lab servers and the four public modelling databases; some models are deposited in more than one database. With two exceptions models were not duplicated in ModelDB and BioModels; the Bhalla and Iyengar [80] model was present in all four public modelling databases, and the Nakano et al. [15] model was found in ModelDB and BioModels. We did not test the functionality or reproducibility of models; only the availability and relative ease of exploration was examined.

### Features of models

We extracted a number of features from each model to highlight their similarities and differences (see Table 3). To quantify the model size, we counted the number of entities that appear in the model. We also extracted information on numbers of dynamic variables per compartment (“Vars/comp.”). Variables are values describing quantities that change in the model. A compartment is defined as a spatial subsection within the model. Since the number of compartments varies with the fineness of the spatial mesh used, the number of variables scales with the number of compartments, but the number of variables per compartment will be a constant, independent of the spatial discretisation used to simulate the model. To provide a measure of model complexity, we used the ratio of thethe number of variables per compartment and the number of entities (“Vars./Comp./Entities”, Table 3).

**Table 2.**
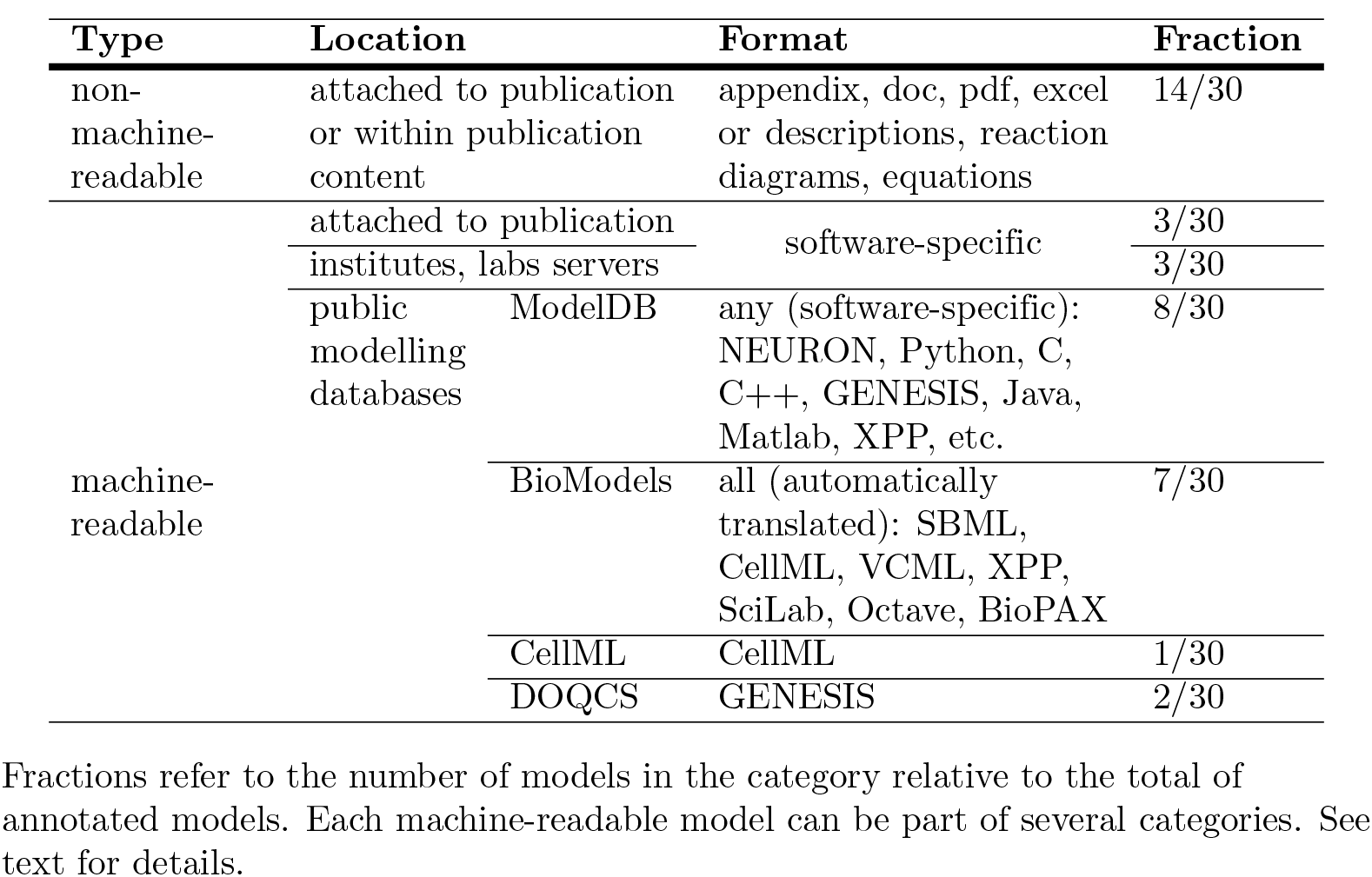
Overview of locations of models and their formats.

For example, in a model of calcium binding to a buffer in a single compartment, there are two entities: calcium (an ion) and the buffer (a protein). There are three variables, namely the concentrations of free calcium, free buffer and calcium-buffer complex. To model diffusion of calcium, buffer and calcium-buffer complex, space could be divided into 100 compartments. The number of variables would then be 300, but the number of variables per compartment would be 3. There would still only be two entities in this model - calcium and the buffer - and the variables per compartment per entity ratio would be 1.5.

A high ratio of variables per compartment to entities reflects a detailed description of a small pathway. For example the model of Byrne et al. [58] - whose stochastic model describes binding of calcium, calmodulin and CaMKII - has 82 variables per compartment and 3 entities, making a ratio of 27.3. The 82 variables correspond to the combinations of calcium bound to the N and C lobes of calmodulin and whether or not these complexes are bound to CaMKII. Dealing with this complexity in the simulation is achieved by using an agent-based Gillespie method (Section “Non-spatial models” in “Biophysical models of synaptic plasticity”). Agent-based simulation also allows the more extreme example of Zeng and Holmes [27], whose model of the Ca^2+^-CaM-CaMKII-PP3 pathway (with calbindin and neurogranin; 6 entities in total) has 14,296,081 possible complexes (i.e. variables), making a ratio of 2,382,680 variables per compartment per entity. At the other end of the spectrum, a low variable to entity ratio indicates larger pathways with each interaction modelled in less detail. For example, the ODE-based model of Bhalla and Iyengar [80], with 44 entities and approximately 100 variables per compartment, has a ratio of 2.3 variables per compartment per entity.

In Table 3 we also indicate the region or cell type the model applies to. Hippocampal CA1 cells are most frequently modelled, followed by striatal MSPNs and cerebellar Purkinje neurons. In some models the location is not specified.

**Table 3.** Summary of models.

| Paper | Vars./comp. | Entities | Vars./comp./<br>Entities | Region |
| --- | --- | --- | --- | --- |
| Antunes and De Schutter (2012) [112] | 103 | 19 | 5.4 | Cereb. Purk. |
| Antunes et al. (2016) [42] |  | 17 |  | Cereb. Purk. |
| Bhalla and Iyengar (1999) [80] | 100 | 42 | 2.4 | Hipp. CA1 Pyr. |
| Byrne et al. (2009) [58] | 82 | 3 | 27.3 | Hipp. CA1 Pyr. |
| Castellani et al. (2001) [90] | 36 | 5 | 7.2 | Cortex** |
| Castellani et al. (2005) [82] | 33 | 13 | 2.5 | Ex. glut. syn.** |
| Graupner and Brunel (2007) [22] | 16 | 5 | 3.2 | Hipp. CA1 Pyr. |
| Gutierrez-Arenas et al. (2014) [18] | 188 | 34 | 5.5 | Striatal MSPN, D1R expressing |
| Hernjak et al. (2005) [26] | 9 | 5 | 1.8 | Cereb. Purk. |
| Khan et al. (2011) [32] | 12 | 1 | 12.0 | Hipp. CA1 Pyr. |
| Kim et al. (2010) [21] | 54 | 18 | 3.0 | Hipp. CA1 Pyr. |
| Kim et al. (2011) [30] | 16 | 17 | 1.0 | Hipp. CA1 Pyr. |
| Kim et al. (2013) [31] | 10 | 18 | 0.6 | Striatal MSPN, mGluR1<br>expressing |
| Kötter (1994) [119] |  | 12 |  | striatal MSPN |
| Kuroda et al. (2001) [111] |  | 20 |  | Cereb. Purk. |
| Li et al. (2012) [33] | 95 | 8 | 11.9 | Generic excitatory spine |
| Mattioni and Le Novère (2013) [34] | 13 | 9 | 1.4 | Striatal MSPN |
| Miller et al. (2005) [49] | 58 | 4 | 14.5 | ** |
| Nair et al. (2015) [17] | 80 | 16 | 5.0 | Striatal MSPN, D1R and D2R<br>expressing* |
| Nakano et al. (2010) [15] | 189 | 28 | 6.8 | Striatal MSPN, D1R expressing |
| Oliveira et al. (2010) [28] | 31 | 9 | 3.4 | HEK293 cells |
| Oliveira et al. (2012) [29] | 113 | 28 | 4.0 | Stratial MSPN |
| Pepke et al. (2010) [20] | 156 | 3 | 52.0 | ** |
| Qi et al. (2010) [19] | 115 | 13 | 8.8 | Stratial MSPN |
| Smolen et al. (2006) [100] | 23 | 9 | 2.6 | Hipp. CA1 Pyr. |
| Smolen et al. (2012) [16] | 14 | 6 | 2.4 | Hipp. CA1 Pyr. |
| Sorokina et al. (2011) [23] | 1,000,000 | 55 | 18,181.8 | Ext. glut. syn. |
| Stefan et al. (2008) [59] | 49 | 3 | 16.3 | ** |
| Zeng and Holmes (2010) [27] | 14,296,081 | 6 | 2,382,680.2 | Hipp. DG |
| Zhabotinsky et al. (2006) [83] | 58 | 11 | 5.3 | Hipp. CA1 Pyr. |

“Paper” refers to the analysed model. “Vars/comp.” is the number of molecular variables per compartment, a measure of the complexity of the model; this was not assessed for all papers. “Entities” is the number of entities in the model, and “Vars./Enties” is the ratio between the number of variables per compartment and the number of entities. This roughly corresponds to the level of detail of the model. “Region” refers to the brain region or cell type where the model is situated (\*\* – no cell specified). Abbreviation: Cereb. Purk., cerebellar Purkinje cell; Ex. glut. syn., excitatory glutamatergic synapse; Hipp. CA1 Pyr., hippocampal CA1 pyramidal cells; Hipp. DG, hippocampal dentate gyrus cell; MSPN, medium spiny projection neuron; \* – denotes that there is more than one model presented in a study and numbers in this table refer to the one with the larger number of “Entities”.

“Paper” refers to the analysed model. “Vars/comp.” is the number of molecular variables per compartment, a measure of the was not assessed for all papers. “Entities” is the number of entities in the model, and een the number of variables per compartment and the number of entities. This roughly il of the model. “Region” refers to the brain region or cell type where the model is situated (** on: Cereb. Purk., cerebellar Purkinje cell; Ex. glut. syn., excitatory glutamatergic synapse; CA1 pyramidal cells; Hipp. DG, hippocampal dentate gyrus cell; MSPN, medium spiny that there is more than one model presented in a study and numbers in this table refer to the “Entities”.

**Table 4.**
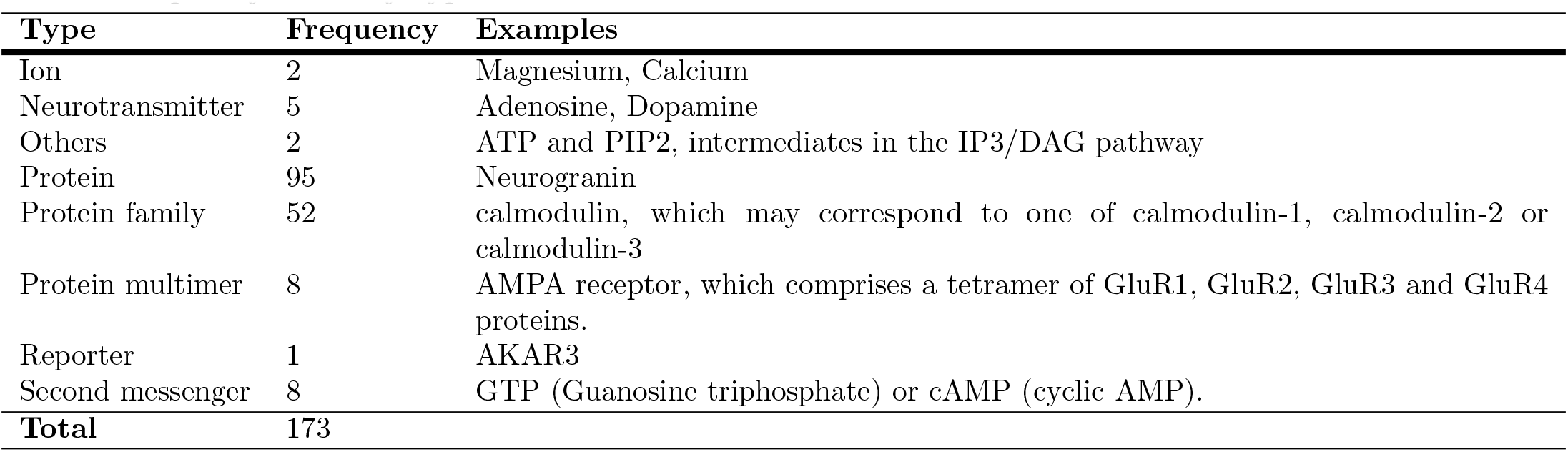
Frequency of entity types found in models.

### Identifying entities in models

To identify the entities in each model, the publication describing the model and, if available, an electronic description of the model were examined by one of the authors. For each entity, we recorded the name used in the model publication and our standard entity identifier. Models do not always specify the entities involved precisely. We discussed ambiguous cases together and erred on the side of not imputing the identity of a protein; for example a “Plasticity related protein” [16] was not mapped to an entity identifier.

We identified 178 distinct entities across the 30 catalogued models (see S1 Table for full list). As well as an identifier, each entity has a long name and a type which can be one of: “ion”, “neurotransmitter”, “others”, “protein”, “protein family”, “protein multimer”, “reporter” or “second messenger”. Table 4 shows how many of each type of entity were identified, and gives examples. The most frequent entity type is “protein”, followed by “protein family” and then “protein multimer”.

The rationale for having three protein types - “proteins”, “protein families” and “protein multimers” - was to allow us to record as precisely as possible what was meant in each computational model. A “protein” is a specific protein e.g. neurogranin, encoded by a specific gene (*NRGN*), so it is unambiguous as to which gene is implied by the model. The same gene may produce multiple isoforms due to gene duplicates or alternate splicing. For example *PRKCZ* produces two isoforms, PKC*ζ* and PKM*ζ* [120]. A “protein multimer” is a multiprotein complex, e.g. an AMPA receptor, which comprises a tetramer of a selection of GluR1, GluR2, GluR3 and GluR4 proteins. In this example, if the model only specified “AMPAR” there would be ambiguity about which of the GluR1-4 subunits are implied by the model. Coding AMPAR as a “protein multimer” allows this ambiguity to be recorded and resolved as desired. A “protein family” is a protein from a family of proteins, e.g. calmodulin, which may correspond to one of calmodulin-1, calmodulin-2 or calmodulin-3. Again, it is not clear which protein is implied by the model, though later we will use information about the synaptic proteome to narrow down the possibilities. “*AKAR3*” is the only entity that was classified as a reporter [17]. The FLIM-AKAR reporter was included in the model to reflect the experimental setup where it is used to measure PKA dynamics. “Ions”, “neurotransmitters” and “second messengers” were assigned to individual classes. They are not proteins, but carry out crucial functions in the cell.

ATP and PIP2, both intermediates in the IP3/DAG pathway were classified as “other”. ATP itself can produce a second messenger and is often referred to as a precursor or “coenzyme”. Similarly, PIP2 is frequently acting as a precursor of a second messenger [31].

The full catalogue of all model entities for all models is shown in matrix form in Fig 4. The models are ordered according to hierarchical clustering (Ward’s 2D method, as implemented in `R’s hclust` function with the `Ward.2D` method). This catalogue is the basis for the rest of the analysis.

**Fig 4.**
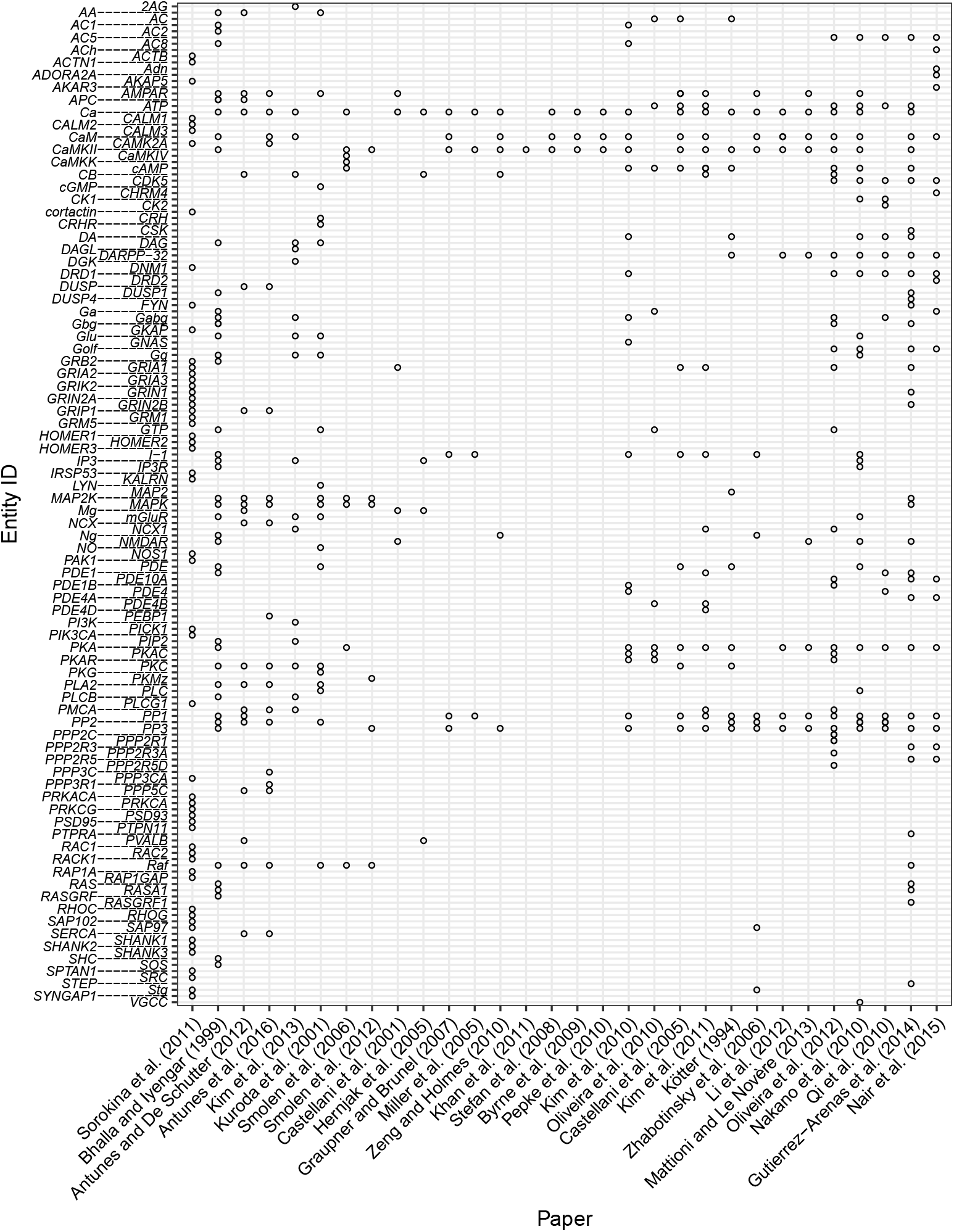
Matrix of entities in models. The occurrence of an entity in a model is indicated by open circles. Entity IDs are staggered for readability.

### Mapping entities to gene identifiers

In order to compare synaptic models with the synaptic proteome, we needed to map each protein entity onto the proteins to which it might correspond. The construction of this mapping is shown in Fig 5. Based on common practice in bioinformatics we decided to use HUGO Gene Nomenclature Committee (HGNC) gene symbols and NCBI Entrez Gene IDs to identify proteins/genes. The one-to-one mapping from HGNC gene symbols to NCBI human Entrez Gene IDs [121] allowed this approach.

As presented in Fig 5, entities of type “protein” were mapped directly to HGNC gene symbols. Entities classified as “protein family” and “protein multimer” required an intermediate mapping step. We searched for ontologies that could be used to identify as many of these entities as possible and map them to HGNC gene symbols. After thorough analysis of available bioinformatic resources (see Methods) we decided to use HGNC gene families to map entities of type “protein family” and “protein multimer” to genes. For each such entity, we tried to identify a corresponding HGNC gene family, and used manual NCBI mapping (see Methods) to check if the genes contained in this family seemed likely to be what was meant in the models. For example, we mapped the entity “Dopamine receptors” (DRD) to the HGNC family “Dopamine receptors”, which contains the genes *DRD1, DRD2, DRD3, DRD4* and *DRD5*. Since this seemed a reasonable set, we accepted the mapping.

For some entities no one HGNC family gave a reasonable set of proteins, but the intersection between two or more families did. For example the genes corresponding to SHANK, by which we mean the family of proteins encoded by *SHANK1, SHANK2* and *SHANK3*, may be selected from the gene families list by choosing all genes that are in the “Ankyrin repeat domain containing” (ANKRD) and “PDZ domain containing” (PDZ) gene families. When we could not find a corresponding HGNC family or a combination of HGNC families, we constructed our own mapping (see Methods). Since “ions”, “neurotransmitters”, “others”, “reporters” and “second messengers” are not proteins, we excluded them from the mapping to gene names.

Once gene families corresponding to 61 “protein families” and “protein multimers” were identified we could map each family or multimer onto a set of genes (S3 Table and S4 Table). 331 unique HGNC gene symbols were identified based on protein families and multimers. The union of this set of symbols with the 96 genes mapped directly from type “protein” in the “full set of HGNC gene symbols in models” dataset. It contains a total of 386 HGNC gene symbols. A number of “protein families” mapped onto the same genes; for example the families PDE and PDE1 both contain *PDE1A* and *PDE1B*.

### Comparison with proteomic data

HGNC families are general gene classes and do not contain information about tissue specificity or expression patterns. To identify proteins found in the synapse, we used a meta-analysis of published proteomic datasets of the presynapse, postsynapse and synaptosome that we are preparing for another publication. The individual references, as of July 2017, can be found in S2 Table.

The synaptosome is the largest data subset and extracted from brain homogenate. The term synaptosome refer to the complete presynaptic terminal including mitochondria, synaptic vesicles and the postsynaptic membrane together with the PSD [122,123]. The PSD is a tightly connected, dense region of the postsynaptic membrane which hosts a number of different receptors and regulatory units. The presynapse and postsynapse are subsets of the synaptosome, and can be separated through experimental steps.

The union of these three datasets, which we refer to as the “synaptic proteome”, comprises 6,706 genes and is based on data obtained from 37 publications and 39 dataseis (data as of July 2017). The extracted proteome was used to filter the “full set of HGNC Gene symbols in models” (see Fig 5 and “Identifying entities in models”). We found that every “protein family” (S3 Table) and “protein multimer” (S4 Table) in our list contains at least one gene overlapping with the synaptic proteome. Genes not expressed in the synapse (“OUT SYNAPSE” in S3 Table and S4 Table) were excluded from further analysis. This filtering step reduces the 331 genes in families to 239 HGNC gene symbols. Together with directly mapped proteins this leaves us with 294 unique HGNC gene symbols describing all mapped genes in models, where families and multimers were screened for the presence in the synapse. From now on we refer to this gene set as “genes in models” (see green box, Fig 5).

**Fig 5.**
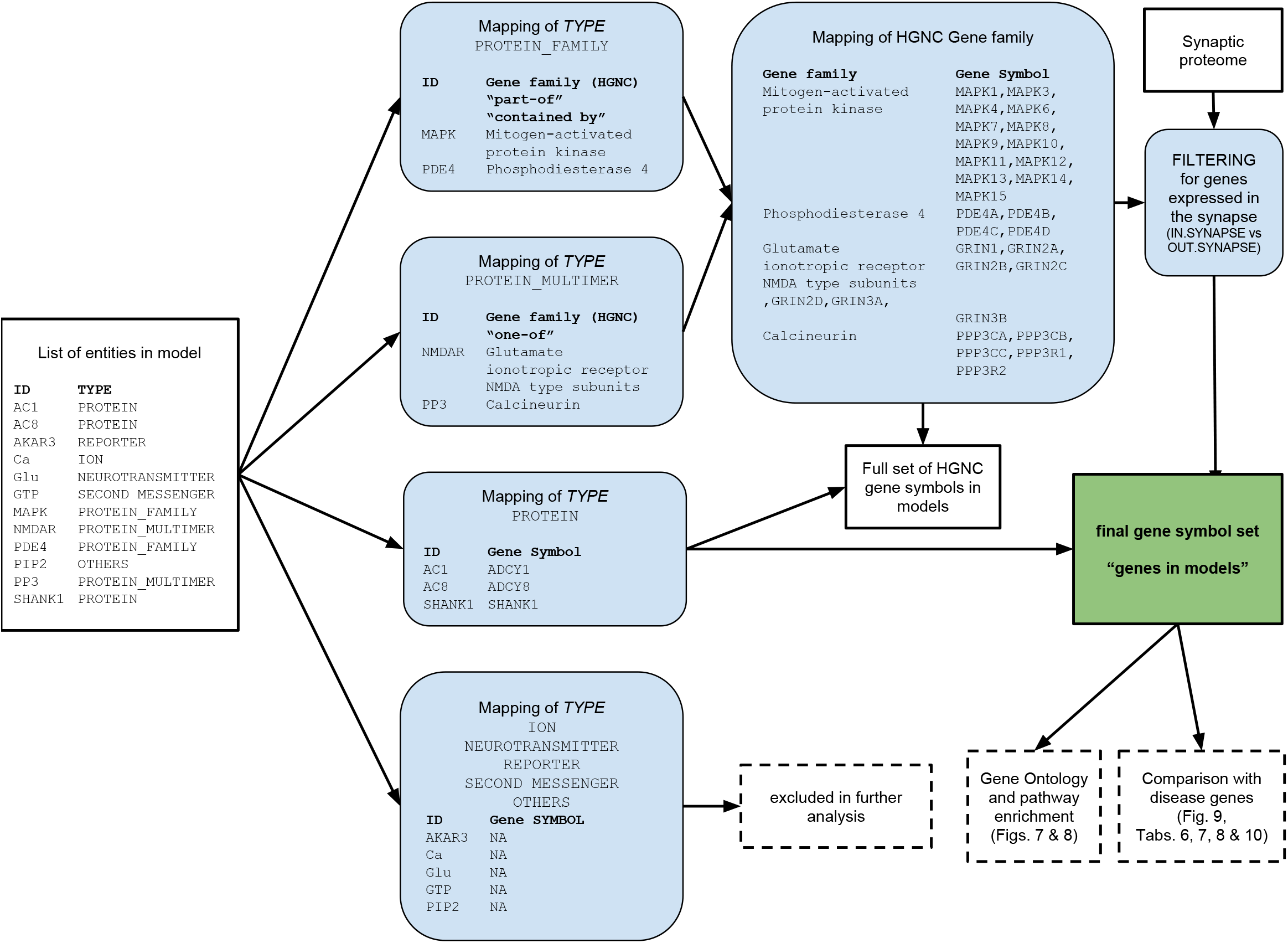
Overview of entity to Gene Symbol mapping process. Sets of data are shown in boxes with black rectangular in boxes with blue backgrounds and curved corners. Dashed lines indicate additional) me is highlighted in a box with green background. Bold font refers to column headers.

The overlap between the final set of “genes in models” and the synaptic proteome, as well as its subsets (presynaptic, postsynaptic, and synaptosome) is visualised in the Venn diagram in Fig 6. It can be seen that 46% of “genes in models” (135 genes) are found in all three synaptic proteome datasets. Significantly lower numbers are expressed in individual sub-datasets. These are 3, 14 and 21 genes for the presynapse, postsynapse and synaptosome respectively (representing 1.0%, 4.7% and 7.1% of genes in models). When disregarding “genes in models” present in the intersection of all three datasets, more modelled genes are found in the postsynapse or synaptosome (143 genes) than the presynapse or synaptosome (27 genes). Thus, postsynaptic genes appear to be the most highly modelled subset. However, relative to the total size of the respective proteomes, only 5.1% of postsynaptic genes (258 “genes in models” out of 5,053 postsynaptic genes) versus 7.6% of presynaptic genes (142 “genes in models” out of 1,867 presynaptic genes) are represented in the models.

**Fig 6.**
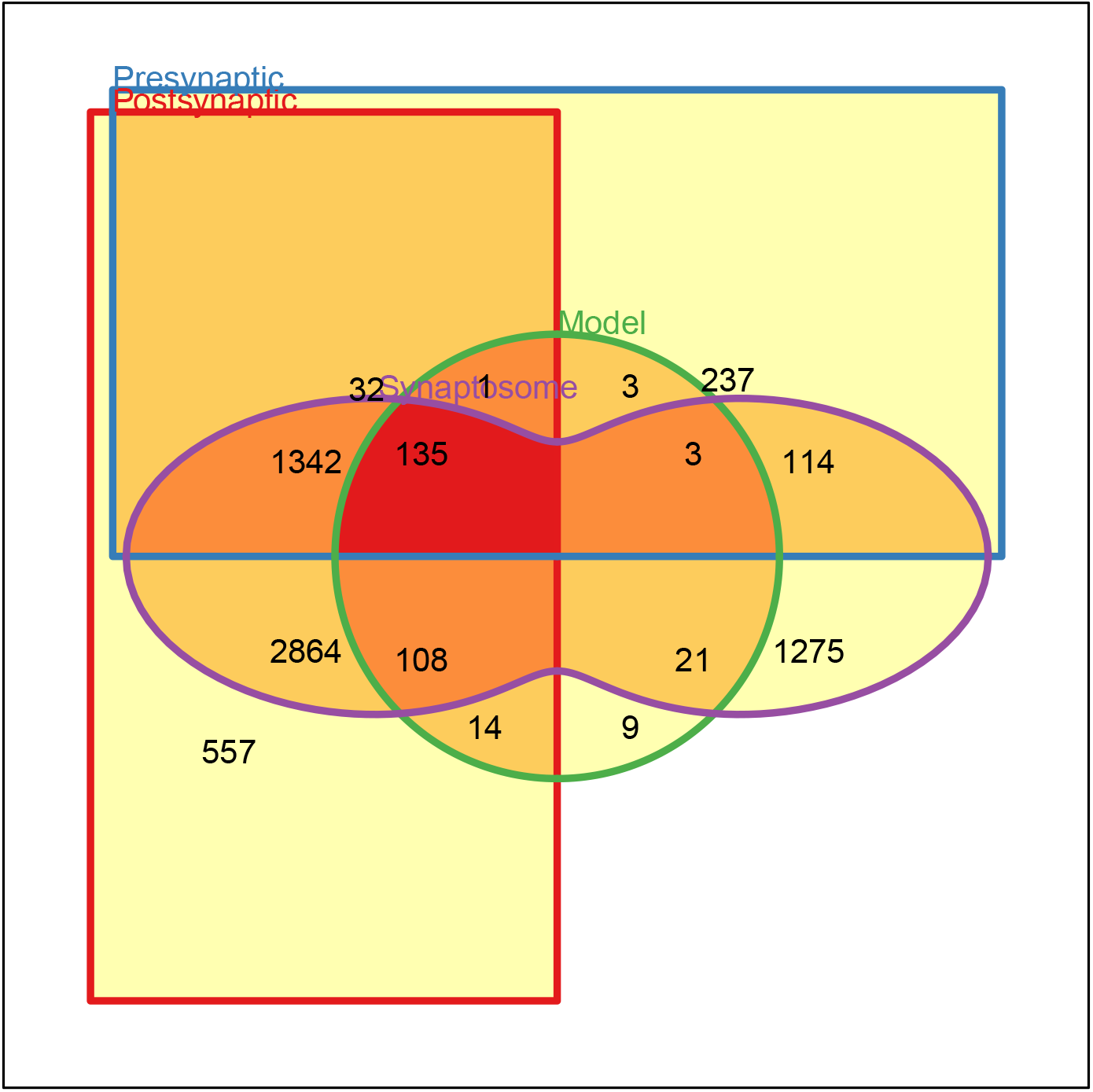
Relationships between the sets of genes in postsynaptic, presynaptic, synaptosome datasets and the sets of genes possibly present in models. Postsynaptic genes in red, presynaptic in blue, the synaptosome in purple and genes in models in green. Numbers refer to the number of genes in each subset and shading shows how many sets a region belongs to (white - none; red - all four). It can be seen that the number of genes in the proteome but not included in models is an order of magnitude bigger than the number of proteins included in models and the proteomic datasets. There are only 9 genes (listed in Table 5) found in models and none of the proteomic datasets.

Nine modelled genes, all of type “protein” are not present in the synaptic proteome datasets (see lower right of the circle in Fig 6). Further investigation shows evidence for all of them being expressed in the synapse (Table 5), so these 9 genes remained in the set of “genes in models”. These cases illustrate how proteomic datasets still seem to be slightly incomplete.

**Table 5.** Proteins in models and not to be found in synaptic datasets.

| Entity ID | Gene | Reason for inclusion |
| --- | --- | --- |
| ADORA2A | *ADORA2A* | Adenosine A2a receptors (A2aR) are expressed with D2R receptors [17] |
| CALM2 | *CALM2* | Unpublished dataset |
| CHRM4 | *CHRM4* | Muscarinic cholinergic receptor shown to be expressed in gonadotropin releasing hormone neurons [124] |
| CRH | *CRH* | Corticotropin-releasing factor, regulating the release of adrenocorticotropin in synapses [125] |
| DRD1 | *DRD1* | D1 subtype of the G-protein coupled dopamine receptor - the most abundant in the central nervous system. [126] confirms the presence in neurons. |
| DRD2 | *DRD2* | D2 subtype of the G-protein coupled dopamine receptor. [126] confirms the presence in neurons. |
| DUSP1 | *DUSP1* | Model specifies that DUSP1 feedback loop occurs in the dendritic shaft, the soma and the nucleus [18] |
| I-1 | *PPP1R1A* | Unpublished dataset |
| PPP2R3A | *PPP2R3A* | Preliminary studies suggest PPP2R3A is present in both cytoplasm and nucleus of cells in the striatum [127]. PPP2R3A mediates Ca<sup>2</sup>-dependent dephosphorylation at Thr-75 of DARPP-32 [127]. |

### Enrichment analysis of modelled genes

After compiling the “genes in models” list, we related it to existing biological knowledge, in the form of gene sets annotated with various biological categories, supplied through a number of databases. Depending on each database’s focus, structured, controlled, and descriptive terms are associated to each gene. As an example for this study, we chose to use the following ontologies: Gene Ontology (GO) [128], REACTOME Pathway Database (REACTOME) [129] and Disease Ontology (DO) [130]. Amongst these GO is the largest and most commonly used ontology, classifying genes within domains including Molecular Function, Biological Process and Cellular Compartment. We also used REACTOME, a free and manually curated database in which genes are tagged with terms representing biochemical reactions and pathways they are involved in. A pathway is composed of one or more reactions or reaction-like events, such as binding, complex formation, transport or polymerisation.

To relate “genes in models” to their associated diseases, we used the DO to provide disease classifications. Multiple sources contain gene disease information. We used annotations retrieved from the GeneRif [131], OMIM [132, 133] and Ensemble Variation [134] databases. Based on annotations in the different ontologies we aimed to identify functionalities shared by the “genes in models”. The `topONTO` package implemented in `R` [135] was used to undertake enrichment analysis (see Methods).

The results are summarised using word clouds to show significantly enriched terms, based on GO annotations, describing Molecular Functions (Fig 7A) and Biological Processs (Fig 7B) for our “genes in models”. It can be seen that a high number of modelled genes are involved in molecular functions such as “G-protein beta/gamma-subunit complex binding”, “GTPase activity”, “calmodulin binding”, “3’,5’-cyclic-AMP phosphodiesterase activity”, “high voltage-gated calcium channel activity”, “signal transducer activity” and “calcium-transporting ATPase activity” amongst others. The most common biological processes are “cellular response to glucagon stimulus”, “platelet activation”, “calcium ion transmembrane transport”, and “activation of protein kinase A activity”.

**Fig 7.**
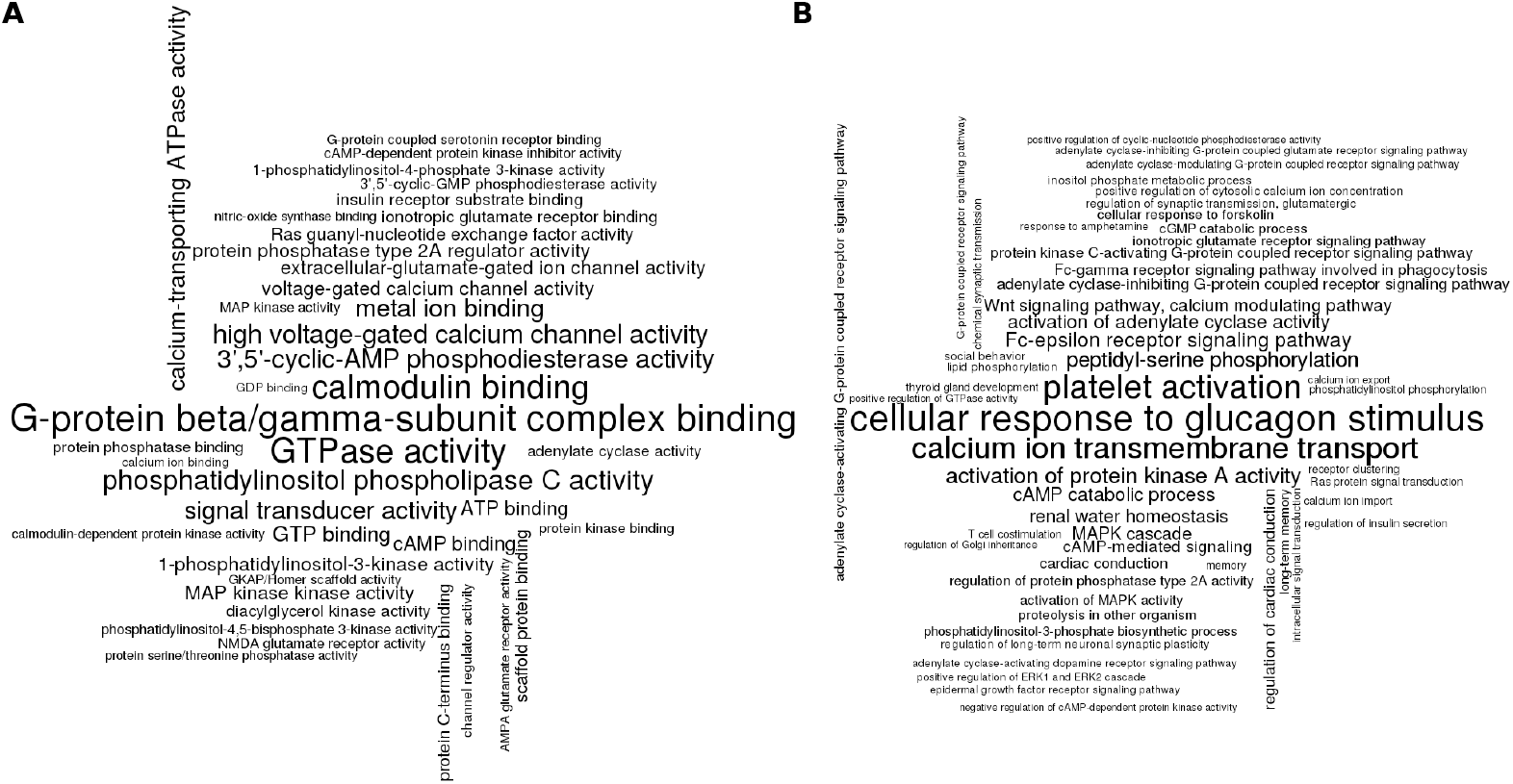
GO enrichment analysis results for “genes in models”. A: Molecular Function ontology terms enriched for “genes in models”. B: Biological Process ontology terms enriched for “genes in models”. The synaptic proteome was used as a background dataset. The list of significant terms was obtained with the Fisher’s exact test and the `elim` algorithm, followed by Benjamini and Yekutieli multiple testing correction. The terms shown in clouds scored less than 0.01 p-value after the correction. Font size is proportional to the term significance.

The identified molecular functions show that genes included in annotated models cover key synaptic processes mainly concentrating around energy production as well as synaptic signalling and information transmission. Identified biological processes are slightly more diverse. Fairly generic processes were identified, showing that the set of modelled genes covers these functions in the synapse. More unique processes appear indicating the synapse specific biological processes described by genes in models.

Fig 8 shows results of the REACTOME enrichment analysis that identified “G alpha (s) signalling events”, “G alpha (z) signalling events” and “DARPP-32 events” as the top enriched pathways. The first two terms are parallel to each other on the pathway hierarchy and have a common parent term of “GPCR downstream signalling”. A comparison of the remaining members of this pathway with the enrichment results shows that they are all significantly enriched in terms of our “genes in models”. The identification of signalling pathways highlights a focus of the analysed models indicating the central role of G-protein signalling.

When considering common diseases amongst “genes in models”, Fig 9A shows a significant enrichment of “schizophrenia” associated genes in the set of “genes in models”, followed by “bipolar disorder”, “Huntington’s disease” and “Alzheimer’s Disease”. The order of results is slightly rearranged when considering the whole cell as a background dataset (Fig 9B). For instance, “Alzheimer’s Disease” becomes more prominent, showing the second highest significance for enrichment in our dataset of interest. On the other hand, “bipolar disorders” drops down the list to the fifth position and “autistic disorder” appears in the results. This shows how different diseases not only affect specific tissues but can affect a larger number of body regions inducing their effect.

**Fig 8.**
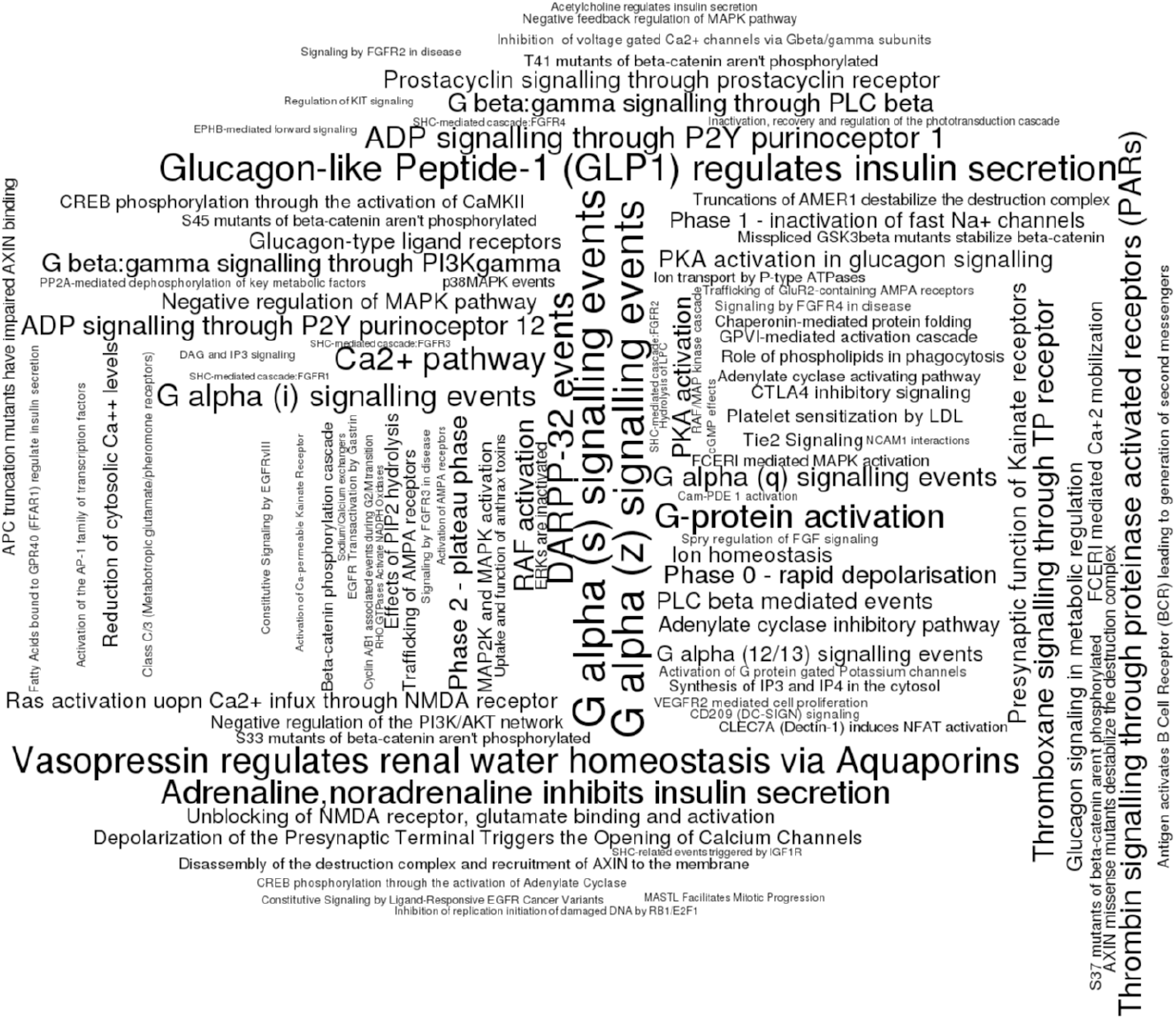
REACTOME enrichment analysis results for “genes in models”. The synaptic proteome was used as background dataset. The list of significant terms was obtained with the Fisher’s exact test and the `elim` algorithm, followed by Benjamini and Yekutieli multiple testing correction. The terms shown in clouds scored less than 0.01 p-value after the correction.

**Fig 9.**
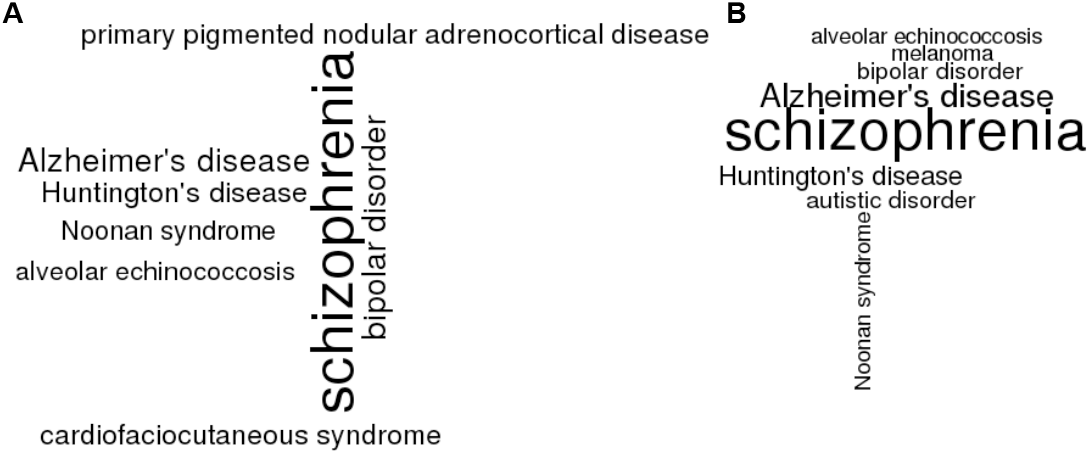
DO enrichment analysis results of “genes in models”. Two background datasets were used: synaptic proteome (A) and all human protein coding genes (B). The list of significant terms was obtained with the Fisher’s exact test and the `elim` algorithm, followed by Benjamini and Yekutieli multiple testing correction. The terms shown in clouds scored less than 0.01 p-value after the correction.

### Modelled genes and their overlap with disease genes

Based on the preceding enrichment analyses we wanted to test for specific associations of modelled genes with disease. Since synapses play a crucial role in signal transduction and are affected in many neurological diseases, these were addressed in more detail. We picked seven representative examples of neurological disorders, 6 of which were based on a list published by the Genes 2 Cognition online initiative: *Attention Deficit Hyperactivity Disorder* (ADHD), *Alzheimer’s Disease* (AD), *Autism, Bipolar Disorder* (BD), *Depression* and *Schizophrenia.* The seventh example was *Parkinson’s Disease* (PD), motivated by our research interests. The list is a representative rather than exhaustive sample of diseases affecting synapses, including diseases of mental health, developmental disorders, as well as diseases of anatomical entity, such as neurodegenerative diseases. Table 6 gives the DO identifiers and short descriptions of each disease.

**Table 6.**
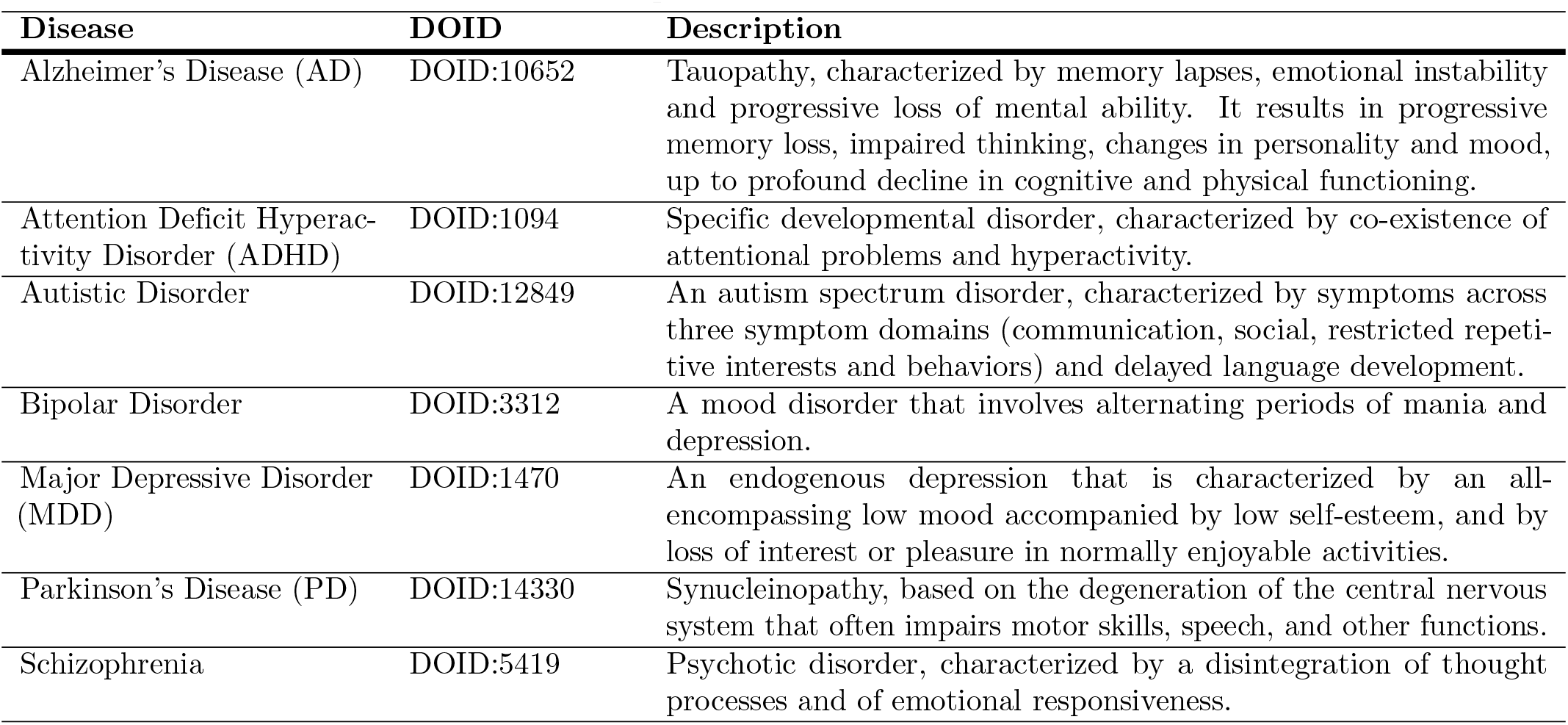
Diseases of Interest and short descriptions.

| Disease | DOID | Description |
| --- | --- | --- |
| Alzheimer's Disease (AD) | DOID:10652 | Tauopathy, characterized by memory lapses, emotional instability and progressive loss of mental ability. It results in progressive memory loss, impaired thinking, changes in personality and mood, up to profound decline in cognitive and physical functioning. |
| Attention Deficit Hyperactivity Disorder (ADHD) | DOID:1094 | Specific developmental disorder, characterized by co-existence of attentional problems and hyperactivity. |
| Autistic Disorder | DOID:12849 | An autism spectrum disorder, characterized by symptoms across three symptom domains (communication, social, restricted repetitive interests and behaviors) and delayed language development. |
| Bipolar Disorder | DOID:3312 | A mood disorder that involves alternating periods of mania and depression. |
| Major Depressive Disorder (MDD) | DOID:1470 | An endogenous depression that is characterized by an all-encompassing low mood accompanied by low self-esteem, and by loss of interest or pleasure in normally enjoyable activities. |
| Parkinson's Disease (PD) | DOID:14330 | Synucleinopathy, based on the degeneration of the central nervous system that often impairs motor skills, speech, and other functions. |
| Schizophrenia | DOID:5419 | Psychotic disorder, characterized by a disintegration of thought processes and of emotional responsiveness. |

Onto Suite Miner [136] was used to obtain all genes linked to the DO IDs from the databases supplying gene-disease association information (GeneRIF, OMIM and EnsemblVariation). The various databases have different approaches to disease-gene annotations. EnsemblVariation relies on genetic mutations (mostly Single Nucleotide Polymorphisms, SNPs), whereas OMIM and GeneRIF contain curated text annotations describing disease-gene associations. These can be queried with text-mining tools and data can be extracted. The different sources were considered individually and jointly. All presented results refer to the full set of disease associated genes irrespective of the original data source. The number of genes linked to each of the diseases can be seen in row: “Disease Genes” in Table 7.

Since not all disease genes are expressed in the synapse, we used the synaptic proteome (Section “Comparison with proteomic data”) to filter the disease associated genes for genes that are expressed in the synapse (see row: “Disease genes in the synapse”, Table 7). Since almost all modelled genes are expressed in the synapse we only present numbers describing the overlap between disease proteins found in the synapse and modelled genes (see row “Disease Genes in Synapse and in Modelled “Genes”, Table 7)

There seem to be large differences in the number range of genes associated with diseases. However, the proportions of genes associated with a disease and expressed in the synapse range between 33% (Bipolar Disorder and Major Depressive Disorder) and 45% (Schizophrenia). When looking at the overlap of modelled genes and disease-associated genes (in the synapse) numbers vary. Schizophrenia seems to have the highest net overlap (92 genes), but also shows the highest number of total associated genes (1844). In total, between 6.1% (Parkinson’s Disease) and 11.8% (Autistic Disorder) of disease genes associated with any of the selected diseases expressed in the synapse appeared in at least one model.

**Table 7.** Overlap of modelled and disease genes.

| Disease | AD | ADHD | Autistic Disorder | Bipolar Disorder | MDD | PD | Schizophrenia |
| --- | --- | --- | --- | --- | --- | --- | --- |
| Disease Genes | 1511 | 665 | 575 | 1140 | 616 | 620 | 1844 |
| Disease Genes in the Synapse | 645 (43%) | 233 (35%) | 255 (44%) | 379 (33%) | 202 (33%) | 262 (42%) | 828 (45%) |
| Disease Genes in Synapse and in modelled Genes | 63 (9.8%) | 20 (8.6%) | 30 (11.8%) | 45 (11.9%) | 23 (11.4%) | 16 (6.1%) | 92 (11.1%) |

Overlap of modelled and disease genes and their presence in the synapse and our modelled gene set. Disease information is based on GeneRif, OMIM and EnsemblVariation database data. “AD” stands for Alzheimer’s Disease, “ADHD” for Attention Deficit Hyperactivity Disorder and “PD” for Parkinson’s Disease. Numbers in brackets refer to the percentages. Percentages in the “Disease Genes in the Synapse” column are relative to the total of “Disease Genes” and “Disease Genes in Synapse and in Modelled Genes” is relative to the number of “Disease Genes in Synapse”.

**Table 8.**
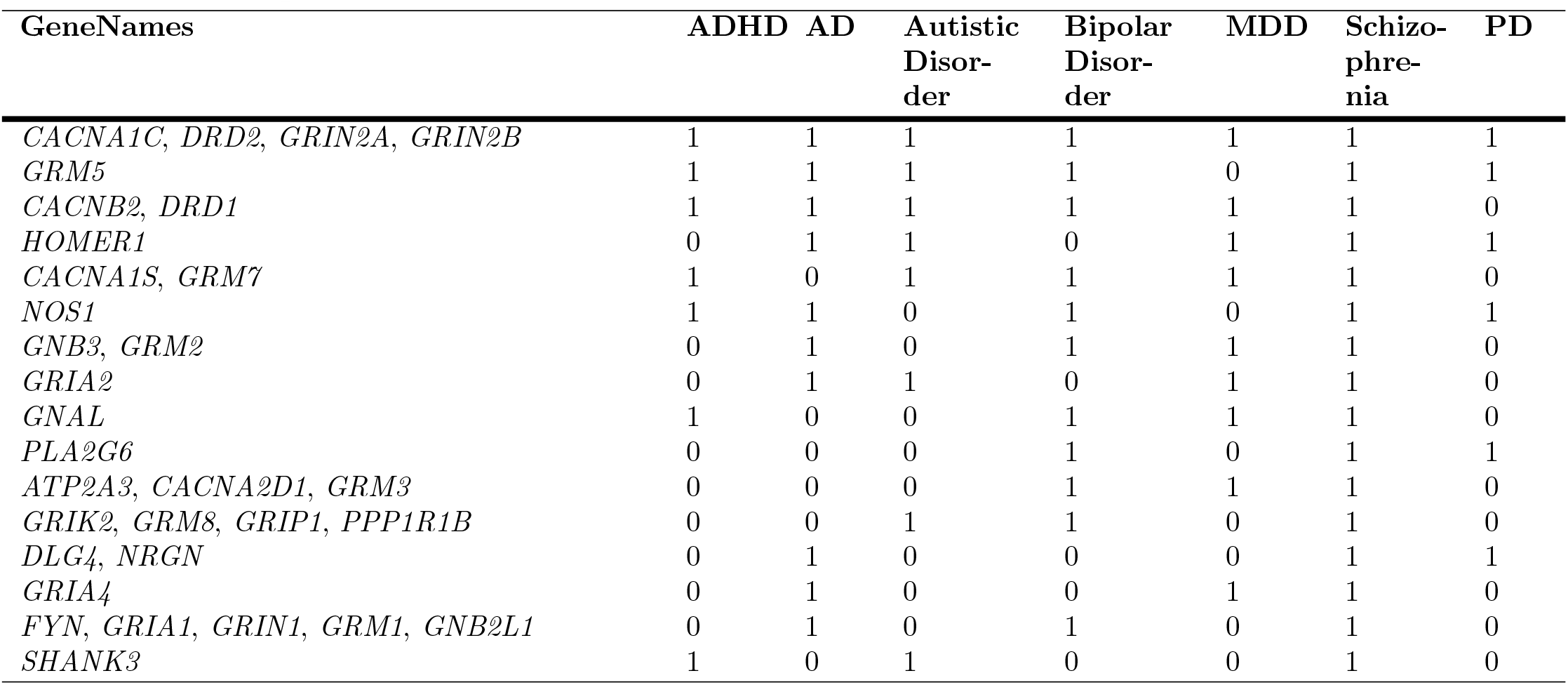
Modelled genes associated with three or more of the selected diseases.

We were also interested in synaptic genes common to a number of diseases. Table 8 shows the 32 synaptic genes linked to three or more of the diseases included in the analysis. Seven genes are associated to six or all seven tested diseases. The top coverage disease associated genes, found in models annotated, include the protein family voltage-dependent calcium channel family *CACNA1C* and *CACNB2* and dopamine D1 and D2 receptors (*DRD1, DRD2*), the inotropic glutamate NMDA receptors, type subunit 2A and 2B (GRIN2A, GRIN2B) as well as the glutamate metabotropic receptor 5 (GRM5). Of the set of modelled genes, 130 (around 50% of the total) are not associated with any of the seven diseases.

In summary, the fraction of genes modelled is relatively small and might indicate that it is challenging to use existing models to make disease predictions. On the other hand the modelled genes can be starting points to extend models to obtain better disease insights, as will be considered later (Approaches to including non-modelled disease genes in models).

### Family trees of entities

Our identification of entities in models makes it possible to query in which models a particular entity is contained. The mapping of entities to genes allows querying models by genes that are, or may be, modelled. It is also desirable to query models by families of molecules. For example Gutierrez-Arenas et al. [18] and Nair et al. [17] include *PDE4A*, whereas Kim et al. [30] and Oliveira et al. [28] include *PDE4B* in their models, and Kim et al. [21] and Qi et al. [19] specify *PDE4*. It would be desirable to be able to search for models containing any of the *PDE4* subfamily of genes.

To enable query by class or family, we determined 29 hierarchical family trees of “proteins”, “protein families” and “protein multimers” implied by the sets of genes corresponding to each (Fig 10). Each “protein family” or “protein multimer” entity is the parent to one or more “proteins” or “protein families”. Each child corresponds to a subset of the proteins in the parent. Tree structures were generated for all “protein multimers” and for “protein families” where a member of that family has been modelled explicitly in at least one of our analysed models. This meant that, for example, PP1 is not represented, since none of its children *PPP1CA, PPP1CB* and *PPP1CC* appear in any model explicitly. Individual proteins appear only if they are part of a family or multimer, and they appear in a model - thus, for example, *GRIA4* and *GRIN3* do not appear. Proteins that do not belong to a family, e.g. *PSD95* (*DLG4*), are not shown.

**Fig 10.**
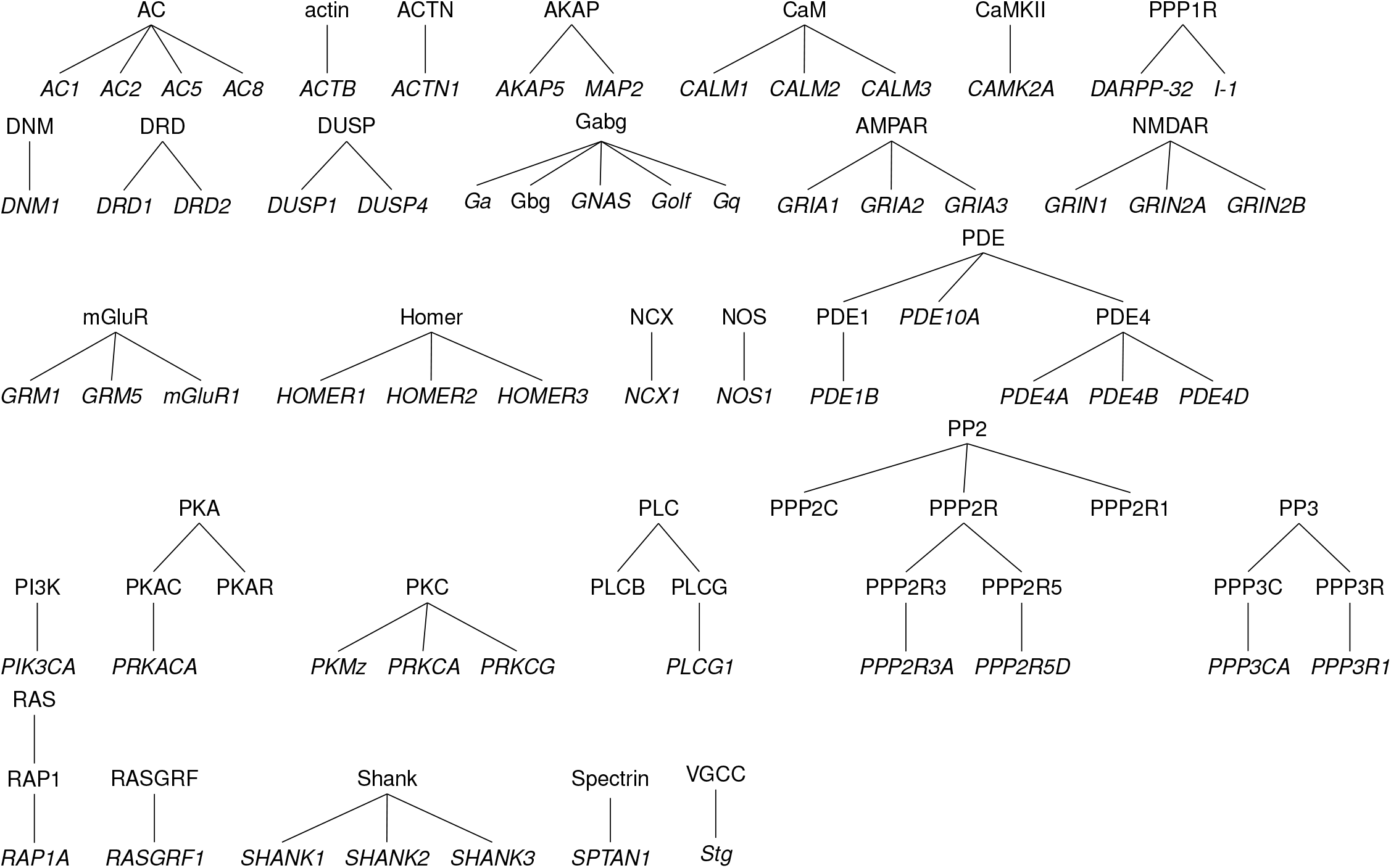
Family trees of “protein families” and “protein multimers”. “Proteins” are shown in italics; “protein rs” in roman. “Proteins” that do not belong to any family are not shown. Only proteins that are specified in models are shown.

Any entity that is part of a family can be mapped to the root node of its tree. Entities that do not belong to a family are implicitly their own root. This mapping of “entities to entity families” (Fig 3) can be applied to the model-entity catalogue (Fig 4) to give the simplified summary mapping of models to 104 family roots shown in Fig 11. This facilitates comparison of entities across models trying to address the differences in model detail between models.

**Fig 11.**
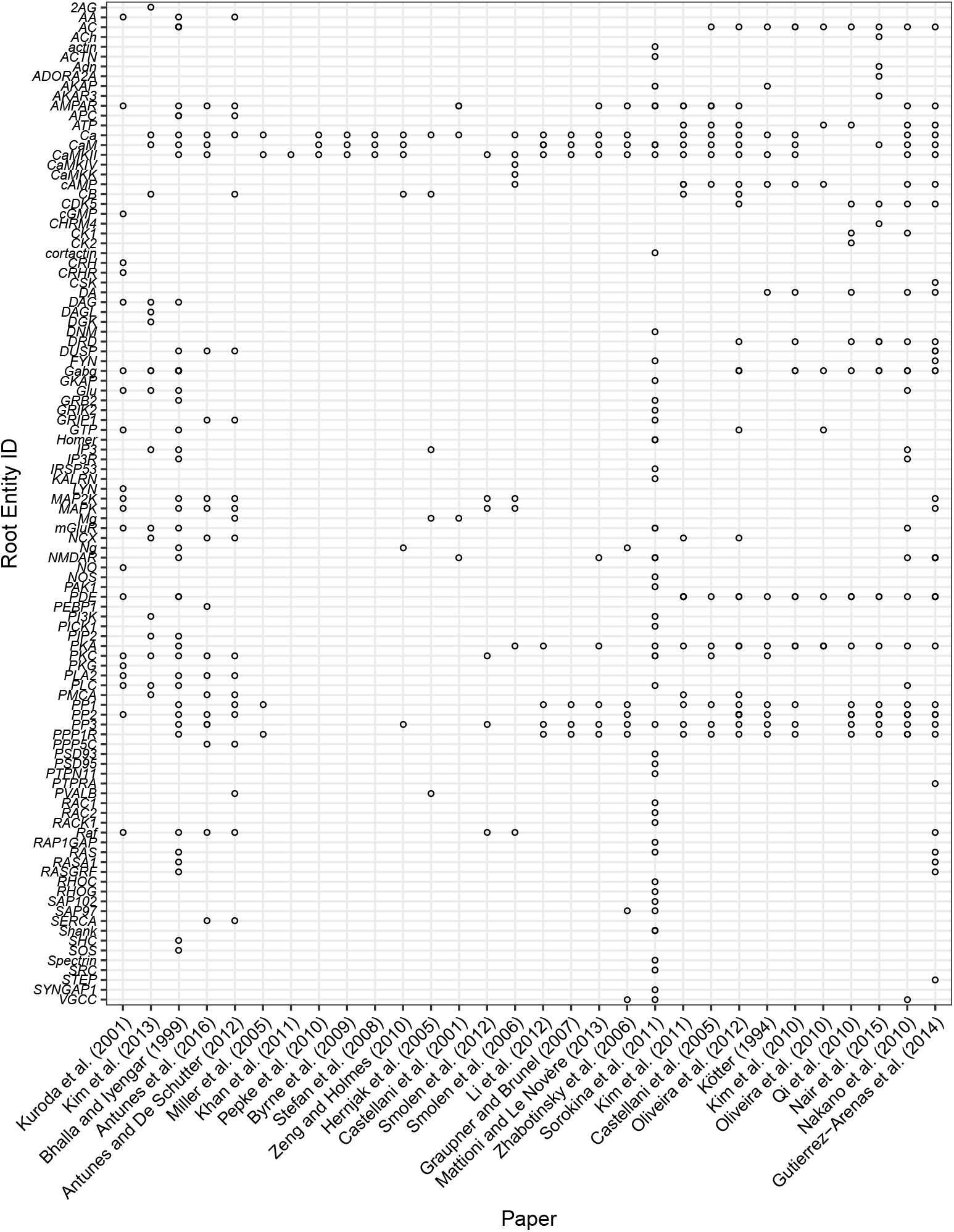
Summary mapping of entities in models. The occurrence of an root entity in a model is indicated by open circles. Lower-level entities are folded into their root entity.

### Frequency of modelling

To give an indication of which are the frequently modelled entities and families of entities, we determined the number of models in which each of the root entities in Fig 12 appears (Table 9). About 50% of root entities appear only in one model. In total, 26 (about 25%) of the entity roots were included in five models or more. The three most frequently modelled entities and families are CaM, CaMKII and Ca, which are included in 18, 22 and 23 out of 30 analysed models respectively. This is due to a number of models focusing specifically on the Ca-CaM-CaMKII pathway or including it as a model part, reflecting its central role in synaptic biology. These top coverage families are followed by families such as PP3 and PP1, PKA and PPP1R, which are also included in the models that include the phosphorylation-dephosphorylation circuit (Section “Models of hippocampal synaptic signalling pathways”). Receptor related families such as AMPAR appear with lower frequency, reflecting the fact that, while crucial for synaptic physiology, not all models include them as a readout mechanism for LTP and LTD. Even though our coverage of models is not complete, it seems likely that cataloguing further models will not change the order much.

**Table 9.**
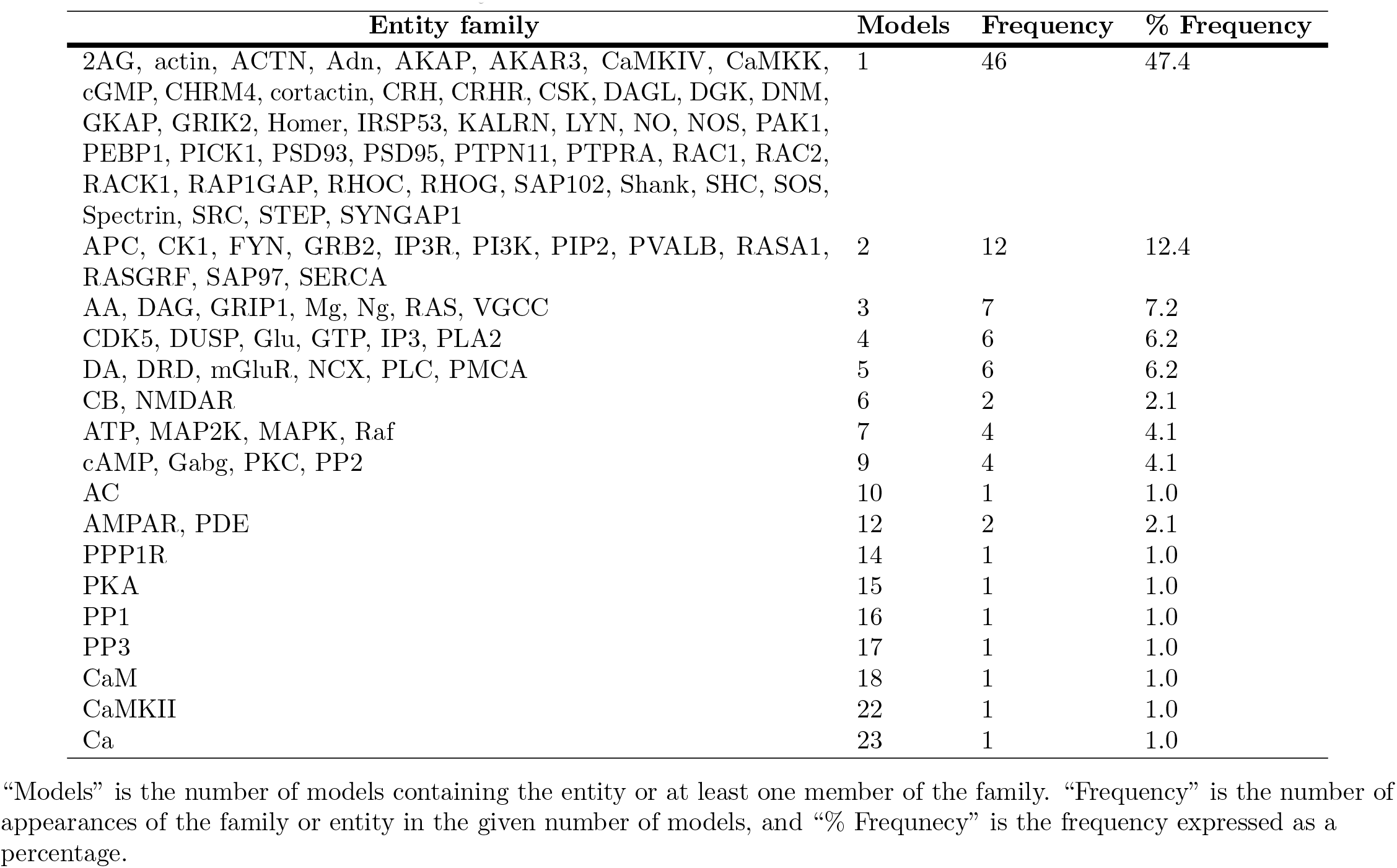
Numbers of entities or entity families found in models.

“Models” is the number of models containing the entity or at least one member of the family. “Frequency” is the number of ntity in the given number of models, and “% Frequnecy” is the frequency expressed as a percentage.

### Comparing models based on their entities

Having annotated the models with entities enabled us to compare models with each other by applying a hierarchical clustering approach to the model-entity root mapping (Fig 11). Ward’s 2D method, as implemented in R’s `hclust` function was used to give the dendrogram shown in Fig 12. We also applied the clustering to the full model-entity matrix (Fig 4), with similar results, though slightly less meaningful groupings.

In Fig 12 similar models cluster together. Three models (Byrne et al. [58], Pepke et al. [20] and Stefan et al. [59]) are clustered together as they all contain the identical set of entities: Ca, CaM and CaMKII. The closely related model of Zeng and Holmes [27] includes CB as well, and the closely related models of Miller et al. [49] and Khan et al. [32] are also centred on CaMKII. The related models of Smolen et al., 2006 [100] and Smolen et al., 2012 [16] feature the MAPK pathway, in addition to CaMKII.

The group of models containing Li et al. [33], Graupner and Brunel [22], Mattioni and Le Novère [34] and Zhabotinsky et al. [83] are all variations on the CaMKII phosphorylation-dephosphorylation circuit, all adding PP1 and PP3 (calcineurin) to the Ca-CaM-CaMKII pathway. All the models so far are hippocampal; Kim et al. [31] is the closest related striatal model to those mentioned. The model of Sorokina et al. [23] is dissimilar to other models, reflecting the large number of entities, particularly scaffolding proteins, which are contained in this model but not in others.

The next cluster contains a sub-cluster of mostly striatal models [15,17-19,21,30], with the exception of Castellani et al. [82], which is one of the few hippocampal models to contain the AC-cAMP-PKA pathway as well as hydrolisation of cAMP to AMP by PDE. The model of Bhalla and Iyengar contains these pathways and many more, accounting for its loose connection with this cluster. In summary, we have shown that models the entity composition can be used to the similarities between models.

**Fig 12.**
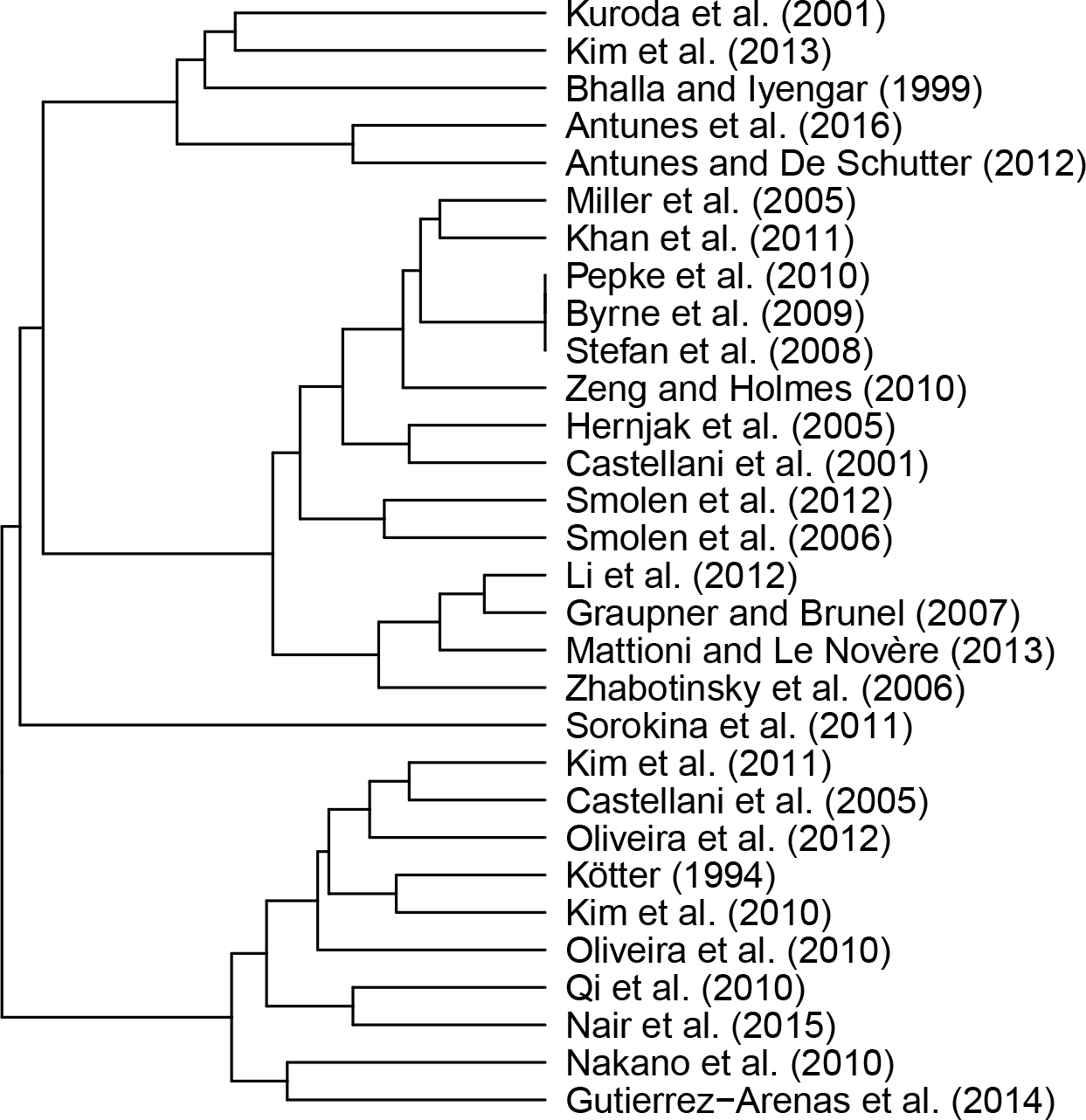
Clustering of model-entity family root matrix. Clustering as implemented in R’s `hclust` function with the `Ward.2D` method.

### Approaches to including non-modelled disease genes in models

Knowing which disease associated genes are included in models helps models with high potential to explain disease impact on the synapse to be identified (“Modelled genes and their overlap with disease genes“). It also allows us to identify disease associated proteins which do not appear in the models we analysed. Of all disease associated genes, 1,248 are found in the synaptic proteome but not in any of the analysed models. Table 10 shows the 32 genes that are associated with 5, 6 or all 7 diseases, and which do not appear in any of the investigated models. Of these, *COMT* and *SLC6A3* are associated with all 7 diseases of interest. Since these genes are associated with all or many studied diseases, they could be of interest when it comes to gaining a better understanding of generic disease dysfunctions.

Supporting the idea that genes implicated in many diseases could be potentially targets for modelling, we identified two genes, *COMT* and *MAOA*, that have been included in metabolic models [137,138]. Functionally, the catechol O-methyltransferase (*COMT*) degrades catechols, such as dopamine, by catalysing their methylation. This methylation results in one of the major degradative pathways of the catecholamine transmitters [139]. Dopamine is included in a number of analysed models [140, 141], and it could be possible to explore what happens in these models if there is an excess of dopamine due to *COMT* malfunction.

**Table 10.**
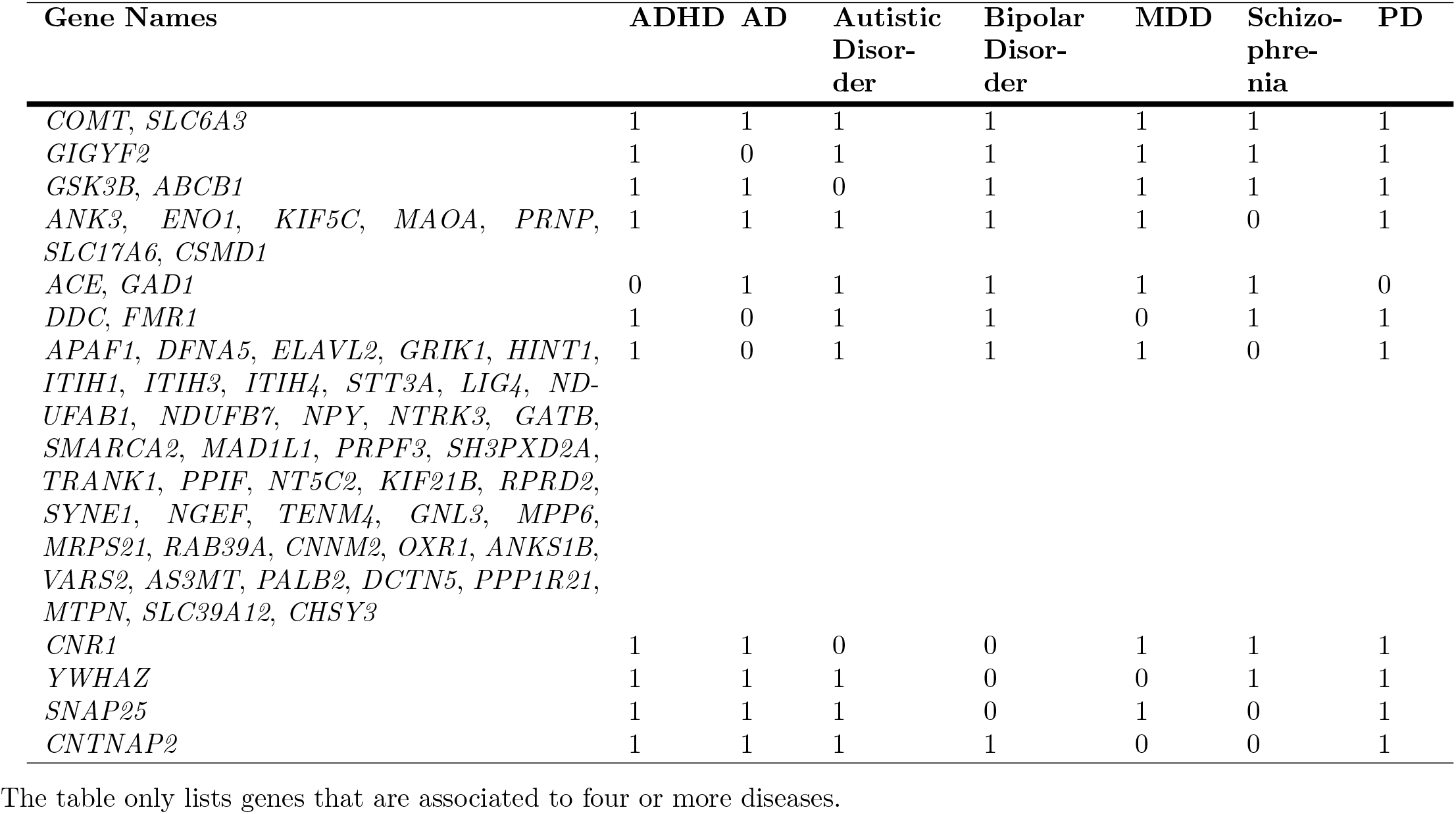
Disease associated genes not appearing in any of the annotated models.

Genes associated will all studied diseases could represent generic disease mechanisms, in which case exploring the role of *COMT* in dopaminergic models would indicate the possible influence of the gene in many diseases. An alternative approach is to consider disease specific genes not appearing in models and associated to only one of the selected diseases. Integrating such proteins into pre-existing models could thus help to gain disease-specific insights. 824 of the disease associated genes are specific to one disease only. To identify genes that can be integrated into existing models, the list of non-modelled disease associated genes was compared with genes in pathways enriched amongst the modelled genes.

For example, all disease genes unique to Schizophrenia were compared with the list of genes in pathways significantly enriched amongst the modelled genes, giving a list of 8 genes, each of which is found in one or more pathways (Table 11). One of these genes is *LAMTOR2*. The LAMTOR2:LAMTOR3 complex binds MAPK components [142],: together with other members of the *MAPK2* and *MAPK* activation pathway, such as: *RAF1, MAPK1, MAPK3* and *MAP2K2*. In this role it contributes to the activation of: the MAPK pathway which has a central role in striatal and cerebellar synapses.

Including the influence of *LAMTOR2* on the activity of MAPK in a pre-existing model could hence help to better understand its role and links to and effects on schizophrenia. Integrating *LAMTOR2* activity in the model could be done mechanistically, or functionally, for example by influencing the MAPK concentration.

**Table 11.** Schizophrenia specific genes not found in models and appearing in pathways that are enriched in annotated models.

| Gene Name | Gene Name (long) | REACTOME pathway | Pathway ID |
| --- | --- | --- | --- |
| *CCK* | cholecystokinin | G alpha (q) signalling events | R-HSA-416476 |
| *LAMTOR2* | late endosomal/lysosomal adaptor, MAPK and MTOR activator 2 | MAP2K and MAPK activation, FCERI mediated MAPK activation, VEGFR2 mediated cell proliferation, RAF/MAP kinase cascade | R-HSA-5674135,<br>R-HSA-2871796,<br>R-HSA-5218921,<br>R-HSA-5673001 |
| *PSMB1* | proteasome subunit beta 1 | FCERI mediated MAPK activation, VEGFR2 mediated cell proliferation, RAF/MAP kinase cascade | R-HSA-2871796,<br>R-HSA-5218921,<br>R-HSA-5673001 |
| *PSMB4* | proteasome subunit beta 4 | FCERI mediated MAPK activation, VEGFR2 mediated cell proliferation, RAF/MAP kinase cascade | R-HSA-2871796,<br>R-HSA-5218921,<br>R-HSA-5673001 |
| *PSMC1* | proteasome 26S subunit and ATPase 1 | FCERI mediated MAPK activation, VEGFR2 mediated cell proliferation, RAF/MAP kinase cascade | R-HSA-2871796,<br>R-HSA-5218921,<br>R-HSA-5673001 |
| *PSMC4* | proteasome 26S subunit and ATPase 4 | FCERI mediated MAPK activation, VEGFR2 mediated cell proliferation, RAF/MAP kinase cascade | R-HSA-2871796,<br>R-HSA-5218921,<br>R-HSA-5673001 |
| *PSMD2* | proteasome 26S subunit and non-ATPase 2 and | FCERI mediated MAPK activation, VEGFR2 mediated cell proliferation, RAF/MAP kinase cascade | R-HSA-2871796,<br>R-HSA-5218921,<br>R-HSA-5673001 |
| *TUBB3* | tubulin beta 3 class III | Chaperonin-mediated protein folding | R-HSA-390466 |

## Discussion

We have developed a catalogue of genes whose corresponding proteins correspond to entities in computational models of synaptic plasticity. To achieve this we developed a new set of standard identifiers for entities in computational models, and mapped those entities corresponding to proteins and protein families onto genes. Although time and lack of machine-readable model descriptions constrained the number of models we could analyse, by selecting models from three brain regions (hippocampus, striatum and cerebellum) we are confident that we have covered the bulk of proteins in models.

We were able to identify 294 genes that could be mapped to entities in computational models. This corresponds to 4.2% of the 6,706 known genes in the synaptic proteome. Enrichment analysis showed that, compared to the set of proteins found in the synapse, the genes in models tended to have more signalling functions, which reflects the focus on signalling pathways in such models. This suggests considerable scope for including new molecules in models. However, models of synapses at the molecular level are already complex and are beset by problems of determining parameters. One strategy to prioritise molecules to add to models is to add those most relevant for disease. Our comparison of the list of genes in models with databases of gene-disease association shows that many disease-associated genes are not currently included in synaptic models, and suggests targets for future modelling.

### Targeting disease-relevant proteins for modelling

The genes in models are more associated with neurological diseases, such as Schizophrenia, Alzheimer’s, Huntington’s disease and bipolar disorder, than randomly selected genes in the synaptic proteome or the whole genome. Nevertheless, depending on the disease, the number of disease-associated genes included in models range between 6% and 12% of the disease-associated genes in the synapse. This suggests that there is considerable potential to include disease-related genes in models. Including these molecules could make these models more useful in helping elucidate disease mechanisms and helping to identify new drug targets.

We identified two un-modelled genes associated with 7 neurological diseases, *COMT* and *MAOA* and we found they have close functional links with existing models. By incorporating pathway enrichment results, we identified *LAMTOR*, a gene uniquely associated with Schizophrenia. *LAMTOR* is linked to the MAP kinase pathway, which features in a number of existing models. This demonstrates the utility of our approach for identifying which proteins to incorporate in existing models so that they can make disease-associated predictions. Further investigation using this approach could indicate other target proteins to add to existing synaptic pathway models to make them more informative about the influence of diseases on the synapse.

### A new ontology for computational neuroscience models

The challenge we faced mapping model entities to genes highlighted a gap between bioinformatics, where each gene is well-defined and has a commonly used identifier, and computational neuroscience, where the elements of models are defined at varying levels of precision: for example they may be proteins, protein families or multimers of proteins. Even within the same model, one element may be specified precisely, for example a particular isoform (PKM*ζ*), and another element may be generic, for example “plasticity related proteins” [16]. From a bioinformatics perspective this may seem offensive, but from the viewpoint of computational neuroscience it is entirely valid: a computational model can be seen as a means to reasoning about a hypothesis; the formulation of the model is the hypothesis and the simulations embody the reasoning that generates the predictions arising from the hypothesis [143]. The modelling process sometimes even requires hypothetical elements, which have no existing identifier. For example, one seminal computational neuroscience model [144] contained hypothetical elements (“gating particles”) that predicted essential features of ion channels function.

The problem of mapping model constituents onto biological entities was noted by the originators of the MIRIAM standard [118]. This standard suggests solving the problem of mapping entities at different levels of abstraction by using a “HasVersion” qualifier to map reactants in models to multiple entities, e.g. to map IP3R to Inositol 1,4,5-triphosphate receoptors type 1, 2 and 3. Most of the models we investigated had not been annotated to MIRIAM standards, and we found it more efficient to define our own ontology containing proteins and protein families. We found that existing ontologies such as UniProt, HGNC gene families [145] and Neurolex [146] were not extensive enough to map proteins specified at different levels of precision (e.g. PDE4A, PDE4) to common families (e.g. PDE), though HGNC gene families covered about half of the protein families we identified.

In the absence of a suitable ontology, we used HGNC gene families and curated other family relationships manually to give a full list of entities (Supporting Information Tables S1) and mappings of proteins to families and multimers in which they occur (Supporting Information Tables S3, S4). These tables form the kernel of an ontology, and we have demonstrated that it can be used to determine the potential genes underlying the proteins in computational models, and to cross-link these genes with expression data. Furthermore, we have demonstrated that the ontology can be used to compare models, for example using hierarchical clustering, and to summarise of how often various protein families have been modelled. By annotating models with identifiers of brain region or neuron type, the set of possible proteins belonging to a model could be narrowed down according to the genes that are expressed in a given region. The same procedure could be used to link the genetic content of synaptic models with other types of data, for example spatial expression data from the Allen Brain atlas. This would make it possible to check that a particular model was valid in the brain region it is supposed to represent, or, conversely, could be used to find brain regions for which a particular model might be valid.

The number of models analysed in this paper was limited by the time it took us to annotate models we had not constructed. While some repositories, such as the curated branch of BioModels, enforce curation of models to MIRIAM standards [118], it would be desirable for all models to be annotated consistently at the time of publication or deposition in a repository. Annotation would be a fairly quick process for authors familiar with the models, and the quality of the information would be higher than if annotated by third parties. Three of the 30 models we investigated were annotated to MIRIAM standards. We did not use the MIRIAM annotations of these models, partly so that our annotation of models was consistent and partly because the MIRIAM standard suggests mapping to external identifiers that are often at a finer level of granularity than we needed to compare models to proteomic data. Were more models curated to MIRIAM standards, it would be worthwhile developing a mapping to our identifiers.

As discussed above, some models are of necessity not precise about which protein is specified. To address this, one option would be for the computational neuroscience and bioinformatics communities to adopt an ontology along the lines of the ones we have generated here. If the ontology were stored in the Interlex dynamic lexicon of biomedical terms, a development of Neurolex [146], it would be straightforward for authors to suggest new terms or relationships. The model metadata could be stored by adding fields to existing repository schema, or our data could be converted to a standalone, API-enabled database.

### Nomenclature

The nomenclature we have used for entities has been decided by the authors. We have been guided by gene names, and some of our choices might be controversial, for example naming PP2B (calcineurin) PP3. Our rationale for using identifiers related to gene names is so there is more consistency between the names of members in a family. For example, in Fig 10, PP3 is the parent of the catalytic and regulatory subunits PPP3C and PPP3R; having PP2B as a parent would not be equally consistent. It would be desirable for the computational neuroscience and bioinformatics communities to agree a common nomenclature.

### New directions in modelling

We have demonstrated the potential of our method of identifying entities in models and mapping them to genes to suggest new, disease-relevant directions for modelling. We believe there is considerable potential for the work to be adopted to suit the needs of the community. Our files are available (S1 File) and suggestions for additions or amendments are welcome. [We will also be making our files available via github.]

More speculatively, despite the challenge of expanding the number and relevant proteins in models of synaptic plasticity, we believe that the time has come to incrementally increase the number of proteins involved in models, especially those involved in disease mechanisms.

## Methods

### Identifying entities in models

The question of what entities mean is outlined in “Analysis of proteins in synaptic models“, subection “Identifying entities in models“. The constituent entities of each model were identified by one of the authors (EMW, KFH or DCS) reading the paper, or extracting elements from a machine-readable representation of the model, for example CellML or Kappa descriptions in the cases of Bhalla and Iyengar [80] and Sorokina et al. [23] respectively. The name used to identify the entity in the model was then mapped to the standardised list of entities that we built up as we looked through the models. In some cases model entities were not specified enough to allow us to map them unambiguously onto a model entity - for example “Plasticity Related Protein” [16]. We did not consider a complex as an entity - for example a Ca-CaM-CaMKII complex would give rise to Ca (ion), CaM (“protein”) and CaMKII (“protein multimer”). In naming our standard entities, we have tried to use names commonly used in models, but for entities that have not appeared in many models we have tended to use the newer standard names that appear in the NCBI or UniProt databases.

### Mapping entities to a unique gene identifier

To obtain a common identifier for all entities we searched for an ontology that could be used to identify our entities, especially “protein families” and “protein multimers”. We considered a number of potential ontologies:

#### The Computational Neuroscience Ontology

(http://bioportal.bioontology.org/ontologies/CNO) This ontology covers the description of the modelling technique (e.g. Integrate-and-fire neurons) rather than the components of the model.

#### HGNC Gene families

(http://www.genenames.org/) The Human Gene Organisation Gene Nomenclature Committee (HGNC) approves unique symbols and names for human genes, and also places genes in families, based on characteristics such as function, homology, domains and phenotype [145]. Placing genes into families is a manual process, often involving specialists who are expert in that family of genes. Often, but not always, genes in the same family have a common root symbol. The process of defining families is ongoing.

#### InterPro protein families

(http://www.ebi.ac.uk/interpro) The InterPro Consortium is a federation amalgamating protein signature databases (Gene3D, Conserved Domain Database, HAMAP, PANTHER, Pfam, PIRSF, PRINTS, ProDom, PROSITE, SMART, SUPERFAMILY, Structure-Function Linkage Database and TIGRFAMs) [147]. Protein signatures are predictive models build on fragments of amino acid sequences that share local features (e.g. conservation at different positions) known to be associated with a function or structure [148]. There are multiple computational approaches that are detecting such patterns and define types of signatures [149]. The similarity in signature matches between proteins is used to define a hierarchy of families.

#### Manual NCBI search

(www.ncbi.nlm.nih.gov/gene/) The National Center for Biotechnology Information (NCBI) provides access to biomedical and genomic information. We used their searchable database of genes, which can be queried with a number of different identifiers.

We intended to map out entities using information supplied by one of these ontologies, but no one source proved sufficient. In InterPro, there are a number of families that correspond exactly to proteins, for example Phospholipase A2 (IPR001211) and Phosphoinositide phospholipase C (IPR001192). However, some proteins, including SOS1 and SOS2, belong to very broad families.

In the HGNC database we identified a relatively large number of our entities that correspond to existing HGNC gene families. For example the HGNC Homer family (short for “Homer scaffolding proteins”) comprises the genes *HOMER1, HOMER2* and *HOMER3* and the genes *PPP3CA, PPP3CB, PPP3CC, PPP3R1* and *PPP3R2* belong to the HGNC PP3 family (short for “Calcineurin”). Other entities do not correspond to a single gene family, but can be extracted from the database by selecting multiple families. For example SHANK, by which we mean the family of proteins encoded by *SHANK1, SHANK2* and *SHANK3* may be selected from the gene families list by: selecting all genes that are in the “Ankyrin repeat domain containing” (ANKRD) and: “PDZ domain containing” (PDZ) gene families. Some of our entities cannot be recovered: by searching for families. For example SOS (by which we mean the proteins encoded by: *SOS1* and *SOS2)* are in both the “Rho guanine nucleotide exchange factors” and: “Pleckstrin homology domain containing” families, but so are 35 other proteins.

We also curated our own mappings by manually querying the NCBI portal by searching for human genes matching a full protein name and a common gene prefix, suffix or infix, if available. For example, Entrez IDs for a “protein family” of Voltage-dependent calcium channel were obtained with the following query: ‘Voltage-dependent calcium channel[All Fields] AND CACN*[All Fields] AND “Homo sapiens”[Organism]’. The top 20 results were considered and only entries with the closest description and gene summary to the search term were extracted.

Although we were not able to map all our entities by relying on only one ontology, we found that HGNC families covered more of our entities than Interpro, so we used this as a basis for developing an ontology to describe the molecular components of computational neuroscience models. We tried to map all entities of type “protein family” and “protein multimer” to HGNC families. Manual NCBI mappings were used to check and verify that HGNC families represented the modelled group of genes.

In situations where we were unable to find a corresponding HGNC family we (1) suggested some protein groups to be added to the list of HGNC families and await approval of the request; (2) we had no choice but to fall back on our manual NCBI mapping. The combination of the above lead us to our final mappings. S3 Table and S4 Table show identified HGNC families as well as the genes belonging to them. The superscript given with the HGNC family name indicates its origin, the official HGNC mapping vs. custom mapping. The columns “IN.SYNAPSE” and “OUT.SYNAPSE” are explained in Section “Comparison with proteomic data“.

### Enrichment Analysis

A commonly used method to find statistically significant commonalities between large gene lists is enrichment analysis, also known as over-representation analysis. Based on information contained in ontological databases, enrichment analysis can show if a set of “genes of interest” contains a significantly high number of genes with the same annotation. This approach allows us to gain a better understanding of underlying common themes in our “genes in models” list.

The underlying principle of such an enrichment analysis is to estimate, for each specific category annotated in the database of interest, if the number of genes in our genes of interest set associated with a certain category is larger than expected by chance. To test this relationship statistically, the hypergeometric distribution or one-tailed Fisher’s exact test is commonly applied. Both are known to be equivalent [150].

The four key numbers required to carry out the statistical calculations are:

1. The number of elements in the full dataset, also considered as the background dataset, N. In our case these are all proteins part of the synaptic proteome.
2. The number of elements n in the subset of the full dataset which is tested for enrichment. This is the number of genes in the “genes in models” list.
3. The number of elements associated to a certain trait in the full dataset, T. It corresponds to the set of genes annotated to any term in one of the databases, e.g. “Schizophrenia”, which describes a disease in the DO database.
4. The subset of n shared by the elements found in T, denoted as t. This refers to the number of genes within a category that are also present in our “genes in models” list.

The probability of encountering the exact number of hits *t* of interest given *N*, *n* and *T* is calculated with the hypergeometric probability *h(t;* N, n, T):

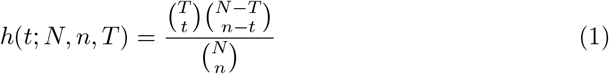

To describe the probability of finding greater than or equal to the number of items of interest *t,* we use the cumulative hypergeometric probability:

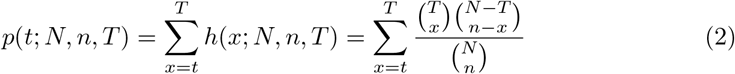

If this probability is less than a criterion (e.g. p < 0.01), the dataset is regarded as enriched [150] for the tested category.

For the analysis, ontology terms for all genes in the background dataset *N* were obtained. Initially two background sets were considered, containing (1) all genes in the genome and (2) all proteins found in the synapse. Since results were quite similar and the focus of this study is on the synaptic region rather than the whole organism, we only present results obtained with the second dataset as the background set of genes.

We analysed all terms that had at least one gene associated to our “genes in models”. For each such term, the *p*-value was calculated, indicating potential enrichment, and then corrected for multiple comparison, using the Benjamini and Yekutieli [151] method. Terms with adjusted *p*-values smaller than 0.01 are presented in the final results.

### topONTO and topGO

Ontologies that supply functional annotation information are organised in a hierarchical structure, with the most generic terms at the top, and the most specific ones at the bottom. The higher the term is located in the hierarchy, the more genes are associated to it as it aggregates all genes from its child terms. Hence, a single gene can be found on different levels of annotation specificity. Depending on the purpose of the analysis it is important to be able to choose the level of retrieved terms.

To retrieve the most specific and refined terms among significantly enriched ones, we used an algorithm proposed by Alexa et al. [152] and implemented for the GO database by the R *topGO* package. Since GO is represented as a Directed Acyclic Graph (DAG), the authors incorporated the underlying GO graph topology in the term scoring approach, removing strong correlations commonly occurring between high level terms. This allows the enrichment of a very generic term to be ignored, and less frequent but more specific and potentially more interesting low level ones to be identified.

Assuming that a child term is potentially more interesting than its more generic ancestors, significance of a term is calculated depending on its child terms. Out of multiple versions implementing this idea, we used the *elim* algorithm paired with Fisher’s exact test. The decision was based on the clear number of comparisons conducted by the algorithm. This number was further used to correct for the false discovery rate.

In the *elim* approach [152], enrichment analysis starts at the bottom of the ontology graph. If a child term is significantly enriched amongst the genes of interest, this influences the number of genes annotated to its ancestor terms. All genes associated to the enriched child term are removed from the ancestor terms leaving most specific ones with the minimal indicated significance.

We discovered that the algorithm leads to more refined results than a set-based enrichment analysis that ignores the ontology structure. Therefore, we were interested in applying a same approach to other gene annotation sets. This can be achieved with the *topOnto* R package [135]. It extends the advantage of the Alexa et al. method to any hierarchically structured dataset. Since both REACTOME and DO satisfy this requirement, we were able to apply the same approach to all chosen annotation sets.

## Supporting information

**S1 File. Data and code.** A zip file containing the data tables, and mapping and analysis code that will reproduce the results in this paper.

**S1 Table. Full list of entities.** List of entities containing the ID, name, type and for proteins, mapping to gene.

**S2 Table. Synaptic Proteome Studies.** List of synaptic proteome publications and respective datasets used in this study.

**S3 Table. Protein family members.** List of entities in distinct protein families - “in” and “out” of the synapse.

**S4 Table. Protein multimer members.** List of entities in distinct protein multimers - “in” and “out” of the synapse.

## References

1. Martin SJ, Grimwood PD, Morris RGM. Synaptic Plasticity and Memory: An Evaluation of the Hypothesis. Annu Rev Neurosci. 2000;23(1):649–711. doi:10.1146/annurev.neuro.23.1.649.

2. Bliss TV, Lømo T. Long-lasting potentiation of synaptic transmission in the dentate area of the anaesthetized rabbit following stimulation of the perforant path. J Physiol (Lond). 1973;232(2):331–356.

3. Lynch GS, Dunwiddie T, Gribkoff V. Heterosynaptic depression: a postsynaptic correlate of long-term depression. Nature. 1977;266:737–739.

4. Abbott LF, Nelson SB. Synaptic plasticity: taming the beast. Nat Neurosci. 2000;3:1178–1183.

5. Nadim F, Bucher D. Neuromodulation of neurons and synapses. Curr Opin Neurobiol. 2014;29:48–56. doi:10.1016/j.conb.2014.05.003.

6. Carlisle HJ, Fink AE, Grant SG, O’Dell TJ. Opposing effects of PSD-93 and PSD-95 on long-term potentiation and spike timing-dependent plasticity. J Physiol (Lond). 2008;586(Pt 24):5885–5900. doi:10.1113/jphysiol.2008.163469.

7. Pocklington AJ, Cumiskey M, Armstrong JD, Grant SGN. The proteomes of neurotransmitter receptor complexes form modular networks with distributed functionality underlying plasticity and behaviour. Mol Syst Biol. 2006;2(1). doi:10.1038/msb4100041.

8. Morrison A, Diesmann M, Gerstner W. Phenomenological models of synaptic plasticity based on spike timing. Biol Cybern. 2008;98(6):459–478. doi:10.1007/s00422-008-0233-1.

9. Manninen T, Hituri K, Kotaleski JHH, Blackwell KT, Linne MLL. Postsynaptic signal transduction models for long-term potentiation and depression. Front Comput Neurosci. 2010;4.

10. Nair AG, Gutierrez-Arenas O, Eriksson O, Jauhiainen A, Blackwell KT, Kotaleski JH. Modeling intracellular signaling underlying striatal function in health and disease. Prog Mol Biol Transl Sci. 2014;123:277–304.

11. Blackwell KT, Jedrzejewska-Szmek J. Molecular mechanisms underlying neuronal synaptic plasticity: systems biology meets computational neuroscience in the wilds of synaptic plasticity. Wiley interdisciplinary reviews Systems biology and medicine. 2013;5(6):717–731.

12. Lassek M, Weingarten J, Volknandt W. The synaptic proteome. Cell Tissue Res. 2015;359(1):255–65. doi:10.1007/s00441-014-1943-4.

13. Bayes A, Collins MO, Croning MD, van de Lagemaat LN, Choudhary JS, Grant SG. Comparative study of human and mouse postsynaptic proteomes finds high compositional conservation and abundance differences for key synaptic proteins. PLoS ONE. 2012;7(10):e46683.

14. Faas GC, Raghavachari S, Lisman JE, Mody I. Calmodulin as a direct detector of Ca^2+^ signals. Nat Neurosci. 2011;14(3):301–304. doi:10.1038/nn.2746.

15. Nakano T, Doi T, Yoshimoto J, Doya K. A kinetic model of dopamine-and calcium-dependent striatal synaptic plasticity. PLoS Comput Biol. 2010;6(2):e1000670.

16. Smolen P, Baxter DA, Byrne JH. Molecular constraints on synaptic tagging and maintenance of long-term potentiation: a predictive model. PLoS Comput Biol. 2012;8(8).

17. Nair AG, Gutierrez-Arenas O, Eriksson O, Vincent P, Hellgren Kotaleski J. Sensing positive versus negative reward signals through adenylyl cyclase-coupled GPCRs in direct and indirect pathway striatal medium spiny neurons. J Neurosci. 2015;35(41):14017–14030.

18. Gutierrez-Arenas O, Eriksson O, Kotaleski JH. Segregation and crosstalk of D1 receptor-mediated activation of ERK in striatal medium spiny neurons upon acute administration of psychostimulants. PLoS Comput Biol. 2014;10(1):e1003445. doi:10.1371/journal.pcbi.1003445.

19. Qi Z, Miller GW, Voit EO. The internal state of medium spiny neurons varies in response to different input signals. BMC Syst Biol. 2010;4(1):1–16. doi:10.1186/1752-0509-4-26.

20. Pepke S, Kinzer-Ursem T, Mihalas S, Kennedy MB. A dynamic model of interactions of Ca^2+^, calmodulin, and catalytic subunits of Ca^2+^/calmodulin-dependent protein kinase II. PLoS Comput Biol. 2010;6(2):e1000675. doi:10.1371/journal.pcbi.1000675.

21. Kim M, Huang T, Abel T, Blackwell KT. Temporal sensitivity of protein kinase A activation in late-phase long term potentiation. PLoS Comput Biol. 2010;6(2):1–14. doi:10.1371/journal.pcbi.1000691.

22. Graupner M, Brunel N. STDP in a bistable synapse model based on CaMKII and associated signaling pathways. PLoS Comput Biol. 2007;3(11):e221.

23. Sorokina O, Sorokin A, Armstrong JD. Towards a quantitative model of the post-synaptic proteome. Mol Biosyst. 2011;7:2813–2823. doi:10.1039/C1MB05152K.

24. Stefan MI, Marshall DP, Le Novere N. Structural analysis and stochastic modelling suggest a mechanism for calmodulin trapping by CaMKII. PLoS ONE. 2012;7(1):e29406. doi:10.1371/journal.pone.0029406.

25. Hepburn I, Chen W, Wils S, De Schutter E. STEPS: efficient simulation of stochastic reaction-diffusion models in realistic morphologies. BMC Syst Biol. 2012;6(1):36.

26. Hernjak N, Slepchenko BM, Fernald K, Fink CC, Fortin D, Moraru II, et al. Modeling and analysis of calcium signaling events leading to long-term depression in cerebellar Purkinje cells. Biophys J. 2005;89(6):3790–3806.

27. Zeng S, Holmes WR. The effect of noise on CaMKII activation in a dendritic spine during LTP induction. J Neurophysiol. 2010;103(4):1798–1808. doi:10.1152/jn.91235.2008.

28. Oliveira RF, Terrin A, Di Benedetto G, Cannon RC, Koh W, Kim M, et al. The Role Of Type 4 Phosphodiesterases in generating microdomains of cAMP: large scale stochastic simulations. PLoS ONE. 2010;5(7):e11725. doi:10.1371/journal.pone.0011725.

29. Oliveira RF, Kim M, Blackwell KT. Subcellular location of PKA controls striatal plasticity: stochastic simulations in spiny dendrites. PLoS Comput Biol. 2012;8(2):e1002383. doi:10.1371/journal.pcbi.1002383.

30. Kim M, Park AJ, Havekes R, Chay A, Guercio LA, Oliveira RF, et al. Colocalization of protein kinase A with adenylyl cyclase enhances protein kinase A activity during induction of long-lasting long-term-potentiation. PLoS Comput Biol. 2011;7(6):e1002084.

31. Kim B, Hawes SL, Gillani F, Wallace LJ, Blackwell KT. Signaling pathways involved in striatal synaptic plasticity are sensitive to temporal pattern and exhibit spatial specificity. PLoS Comput Biol. 2013;9(3):e1002953. doi:10.1371/journal.pcbi.1002953.

32. Khan S, Zou Y, Amjad A, Gardezi A, Smith CL, Winters C, et al. Sequestration of CaMKII in dendritic spines in silico. J Comput Neurosci. 2011;31(3):581–594.

33. Li L, Stefan MI, Le Novere N. Calcium input frequency, duration and amplitude differentially modulate the relative activation of calcineurin and CaMKII. PLoS ONE. 2012;7(9):e43810+. doi:10.1371/journal.pone.0043810.

34. Mattioni M, Le Novere N. Integration of biochemical and electrical signaling - multiscale model of the medium spiny neuron of the striatum. PLoS ONE. 2013;8(7):e66811. doi:10.1371/journal.pone.0066811.

35. Lisman JE. A mechanism for memory storage insensitive to molecular turnover: a bistable autophosphorylating kinase. Proc Natl Acad Sci USA. 1985;82(9):3055–3057.

36. Kuret J, Schulman H. Mechanism of autophosphorylation of the multifunctional Ca2+/calmodulin-dependent protein kinase. J Biol Chem. 1985;260(10):6427–6433.

37. Miller SG, Kennedy MB. Distinct forebrain and cerebellar isozymes of type II Ca2+/calmodulin-dependent protein kinase associate differently with the postsynaptic density fraction. J Biol Chem. 1985;260(15):9039–9046.

38. Lisman JE, Goldring MA. Feasibility of long-term storage of graded information by the Ca2+/calmodulin-dependent protein kinase molecules of the postsynaptic density. Proc Natl Acad Sci USA. 1988;85(14):5320–5324.

39. Zhabotinsky AM. Bistability in the Ca2+/Calmodulin-dependent protein kinase-phosphatase system. Biophys J. 2000;79(5):2211–2221. doi:http://dx.doi.org/10.1016/S0006-3495(00)76469-1.

40. Petersen JD, Chen X, Vinade L, Dosemeci A, Lisman JE, Reese TS. Distribution of postsynaptic density (PSD)-95 and Ca^2+^/calmodulin-dependent protein kinase II at the PSD. J Neurosci. 2003;23(35):11270–8.

41. Gillespie DT. A general method for numerically simulating the stochastic time evolution of coupled chemical reactions. J Comput Phys. 1976;22(4):403–434. doi:http://dx.doi.org/10.1016/0021-9991(76)90041-3.

42. Antunes G, Roque AC, Simoes-de Souza FM. Stochastic induction of long-term potentiation and long-term depression. Sci Rep. 2016;6:30899. doi:10.1038/srep30899.

43. Gaertner TR, Kolodziej SJ, Wang D, Kobayashi R, Koomen JM, Stoops JK, et al. Comparative analyses of the three-dimensional structures and enzymatic properties of alpha, beta, gamma and delta isoforms of Ca^2+^-calmodulin-dependent protein kinase II. J Biol Chem. 2004;279(13):12484–12494. doi:10.1074/jbc.M313597200.

44. Chao LH, Stratton MM, Lee IH, Rosenberg OS, Levitz J, Mandell DJ, et al. A mechanism for tunable autoinhibition in the structure of a human Ca2+/calmodulin-dependent kinase II holoenzyme. Cell. 2011;146(5):732–45. doi:10.1016/j.cell.2011.07.038.

45. Lisman J, Yasuda R, Raghavachari S. Mechanisms of CaMKII action in long-term potentiation. Nat Rev Neurosci. 2012;13(3):169–182. doi:10.1038/nrn3192.

46. Weisstein EW. Necklace; 2017. From MathWorld-A Wolfram Web Resource. Available from: http://mathworld.wolfram.com/Necklace.html.

47. Coomber CJ. Site-selective autophosphorylation of Ca^2+^/calmodulin-dependent protein kinase II as a synaptic encoding mechanism. Neural Comput. 1998;10:1653–1678.

48. Kubota Y, Bower JM. Transient versus asymptotic dynamics of CaM Kinase II: possible roles of phosphatase. J Comput Neurosci. 2001;11(3):263–279. doi:10.1023/A:1013727331979.

49. Miller P, Zhabotinsky AM, Lisman JE, Wang XJJ. The stability of a stochastic CaMKII switch: dependence on the number of enzyme molecules and protein turnover. PLoS Biol. 2005;3(4):e107+. doi:10.1371/journal.pbio.0030107.

50. Stefan MI, Bartol TM, Sejnowski TJ, Kennedy MB. Multi-state modeling of biomolecules. PLoS Comput Biol. 2014;10(9):e1003844. doi:10.1371/journal.pcbi.1003844.

51. Michelson S, Schulman H. CaM kinase: a model for its activation and dynamics. J Theor Biol. 1994;171(3):281–290. doi:10.1006/jtbi.1994.1231.

52. Gillespie D. Approximate accelerated stochastic simulation of chemically reacting systems. J Chem Phys. 2001;115:1716–1733.

53. Holmes WR. Models of calmodulin trapping and CaM kinase II activation in a dendritic spine. J Comput Neurosci. 2000;8(1):65–85.

54. Le Novere N, Shimizu TS. STOCHSIM: modelling of stochastic biomolecular processes. Bionformatics. 2001;17(6):575–6.

55. Danos V, Feret J, Fontana W, Krivine J. Scalable Simulation of Cellular Signaling Networks. In: Shao Z, editor. Programming Languages and Systems. vol. 4807 of Lecture Notes in Computer Science. Berlin, Heidelberg: Springer; 2007. p. 139–157. Available from: http://dx.doi.org/10.1007/978-3-540-76637-7_10.

56. Faeder JR, Blinov ML, Hlavacek WS. Rule-Based Modeling of Biochemical Systems with BioNetGen. In: Maly IV, editor. Systems Biology. vol. 500 of Methods in Molecular Biology. Humana Press; 2009. p. 113–167. Available from: http://dx.doi.org/10.1007/978-1-59745-525-1_5.

57. Sneddon MW, Faeder JR, Emonet T. Efficient modeling, simulation and coarse-graining of biological complexity with NFsim. Nat Methods. 2011;8(2):177–183. doi:10.1038/nmeth.1546.

58. Byrne MJ, Putkey JA, Waxham NN, Kubota Y. Dissecting cooperative calmodulin binding to CaM kinase II: a detailed stochastic model. J Comput Neurosci. 2009;27(3):621–638. doi:10.1007/s10827-009-0173-3.

59. Stefan MI, Edelstein SJ, Le Novere N. An allosteric model of calmodulin explains differential activation of PP2B and CaMKII. Proc Natl Acad Sci USA. 2008;105(31):10768–10773. doi:10.1073/pnas.0804672105.

60. Crouch TH, Klee CB. Positive cooperative binding of calcium to bovine brain calmodulin. Biochemistry. 1980;19(16):3692–8.

61. Colquhoun D, Lape R. Perspectives on: conformational coupling in ion channels: allosteric coupling in ligand-gated ion channels. J Gen Physiol. 2012;140(6):599–612. doi:10.1085/jgp.201210844.

62. Gaertner TR, Putkey JA, Waxham MN. RC3/Neurogranin and Ca^2+^/calmodulin-dependent protein kinase II produce opposing effects on the affinity of calmodulin for calcium. J Biol Chem. 2004;279(38):39374–82. doi:10.1074/jbc.M405352200.

63. Hanson PI, Meyer T, Stryer L, Schulman H. Dual role of calmodulin in autophosphorylation of multifunctional cam kinase may underlie decoding of calcium signals. Neuron. 1994;12(5):943–956. doi:10.1016/0896-6273(94)90306-9.

64. Dupont G, Houart G, De Koninck P. Sensitivity of CaM kinase II to the frequency of Ca2+ oscillations: a simple model. Cell Calcium. 2003;34(6):485–497.

65. Gamble E, Koch C. The dynamics of free calcium in dendritic spines in response to repetitive synaptic input. Science. 1987;236(4806):1311–5.

66. Holmes WR, Levy WB. Insights into associative long-term potentiation from computational models of NMDA receptor-mediated calcium influx and intracellular calcium concentration changes. J Neurophysiol. 1990;63(5):1148–1168.

67. Zador A, Koch C, Brown TH. Biophysical Model of a Hebbian synapse. Proc Natl Acad Sci USA. 1990;87:6718–6722.

68. McDougal RA, Hines ML, Lytton WW. Reaction-diffusion in the NEURON simulator. Front Neuroinform. 2013;7(28). doi:10.3389/fninf.2013.00028.

69. Hepburn I, Chen W, Wils S, De Schutter E. STEPS: efficient simulation of stochastic reaction-diffusion models in realistic morphologies. BMC Systems Biology. 2012;6(1):36. doi:10.1186/1752-0509-6-36.

70. Gillespie DT. Stochastic simulation of chemical kinetics. Annu Rev Phys Chem. 2007;58:35–55. doi:10.1146/annurev.physchem.58.032806.104637.

71. Sorokina O, Sorokin A, Armstrong JD, Danos V. A simulator for spatially extended kappa models. Bionformatics. 2013; p. 3105–3106.

72. Kerr RA, Bartol TM, Kaminsky B, Dittrich M, Chang JC, Baden SB, et al. Fast Monte Carlo simulation methods for biological reaction-diffusion systems in solution and on surfaces. SIAM J Sci Comput. 2008;30(6):3126. doi:10.1137/070692017.

73. Andrews SS. Smoldyn: particle-based simulation with rule-based modeling, improved molecular interaction and a library interface. Bionformatics. 2017;33(5):710–717. doi:10.1093/bioinformatics/btw700.

74. Franks KM, Bartol TM Jr, Sejnowski TJ. A Monte Carlo model reveals independent signaling at central glutamatergic synapses. Biophys J. 2002;83(5):2333–48. doi:10.1016/S0006-3495(02)75248-X.

75. Franks KM, Bartol TM, Sejnowski TJ. An {MCell} model of calcium dynamics and frequency-dependence of calmodulin activation in dendritic spines. Neurocomputing. 2001;38-40:9–16. doi:https://doi.org/10.1016/S0925-2312(01)00415-5.

76. Keller DX, Franks KM, Bartol TM Jr, Sejnowski TJ. Calmodulin activation by calcium transients in the postsynaptic density of dendritic spines. PLoS ONE. 2008;3(4):e2045. doi:10.1371/journal.pone.0002045.

77. Weinan E, Lu J. Multiscale modeling. Scholarpedia. 2011;6(10):11527.

78. Sterratt DC, Sorokina O, Armstrong JD. Integration of Rule-Based Models and Compartmental Models of Neurons. In: Maler O, Halász A, Dang T, Piazza C, editors. Hybrid Systems Biology: Second International Workshop, HSB 2013, Taormina, Italy, September 2, 2013 and Third International Workshop, HSB 2014, Vienna, Austria, July 23-24, 2014, Revised Selected Papers. vol. 7699 of LNBI. Cham: Springer International Publishing; 2015. p. 143–158. Available from: http://dx.doi.org/10.1007/978-3-319-27656-4_9.

79. Lisman J. A mechanism for the Hebb and the anti-Hebb processes underlying learning and memory. Proc Natl Acad Sci USA. 1989;86(23):9574–9578.

80. Bhalla US, Iyengar R. Emergent Properties of Networks of Biological Signalling Pathways. Science. 1999;283:381–387.

81. Ajay SM, Bhalla US. A role for ERKII in synaptic pattern selectivity on the time-scale of minutes. Eur J Neurosci. 2004;20(10):2671–2680. doi:10.1111/j.1460-9568.2004.03725.x.

82. Castellani GC, Quinlan EM, Bersani F, Cooper LN, Shouval HZ. A model of bidirectional synaptic plasticity: from signaling network to channel conductance. Learning and MemoryLearn Memory http://wwwlearnmemorg/. 2005;12(4):423–432.

83. Zhabotinsky AM, Camp RN, Epstein IR, Lisman JE. Role of the Neurogranin Concentrated in Spines in the Induction of Long-Term Potentiation. J Neurosci. 2006;26(28):7337–7347. doi:10.1523/jneurosci.0729-06.2006.

84. Urakubo H, Honda M, Froemke RC, Kuroda S. Requirement of an allosteric kinetics of NMDA receptors for spike timing-dependent plasticity. J Neurosci. 2008;28(13):3310–3323. doi:10.1523/jneurosci.0303-08.2008.

85. D’Alcantara P, Schiffmann SN, Swillens S. Bidirectional synaptic plasticity as a consequence of interdependent Ca2+-controlled phosphorylation and dephosphorylation pathways. Eur J Neurosci. 2003;17(12):2521–2528.

86. Bear MF. Bidirectional synaptic plasticity: from theory to reality. Philos Trans R Soc Lond, B, Biol Sci. 2003;358(1432):649–655.

87. Barria A, Muller D, Derkach V, Griffith LC, Soderling TR. Regulatory phosphorylation of AMPA-type glutamate receptors by CaM-KII during long-term potentiation. Science. 1997;276(5321):2042–2045.

88. Lee HK, Barbarosie M, Kameyama K, Bear MF, Huganir RL. Regulation of distinct AMPA receptor phosphorylation sites during bidirectional synaptic plasticity. Nature. 2000;405(6789):955–959. doi:10.1038/35016089.

89. Lee HK, Kameyama K, Huganir RL, Bear MF. NMDA Induces Long-Term Synaptic Depression and Dephosphorylation of the GluR1 Subunit of AMPA Receptors in Hippocampus. Neuron. 1998;21(5):1151–1162. doi:10.1016/s0896-6273(00)80632-7.

90. Castellani GC, Quinlan EM, Cooper LN, Shouval HZ. A biophysical model of bidirectional synaptic plasticity: Dependence of AMPA and NMDA receptors. Proc Natl Acad Sci USA. 2001;98:12772–12777.

91. Bienenstock EL, Cooper LN, Munro PW. Theory for the development of neuron selectivity: orientation specificity and binocular interaction in visual cortex. J Neurosci. 1982;2:32–48.

92. Liischer C, Xia H, Beattie EC, Carroll RC, von Zastrow M, Malenka RC, et al. Role of AMPA Receptor Cycling in Synaptic Transmission and Plasticity. Neuron. 1999;24(3):649–658. doi:10.1016/s0896-6273(00)81119-8.

93. Choquet D, Triller A. The dynamic synapse. Neuron. 2013;80(3):691–703.

94. Opazo P, Choquet D. A three-step model for the synaptic recruitment of AMPA receptors. Mol Cell Neurosci. 2011;46(1):1–8.

95. Esteban JA, Shi SHH, Wilson C, Nuriya M, Huganir RL, Malinow R. PKA phosphorylation of AMPA receptor subunits controls synaptic trafficking underlying plasticity. Nat Neurosci. 2003;6(2):136–143. doi:10.1038/nn997.

96. Granger AJ, Nicoll RA. Expression mechanisms underlying long-term potentiation: a postsynaptic view, 10 years on. Philos Trans R Soc Lond, B, Biol Sci. 2014;369(1633):20130136+. doi:10.1098/rstb.2013.0136.

97. Migliore M, Hoffman DA, Magee JC, Johnston D. Role of an A-Type K+ Conductance in the Back-Propagation of Action Potentials in the Dendrites of Hippocampal Pyramidal Neurons. J Comput Neurosci. 1999;7:5–15.

98. Poirazi P, Brannon T, Mel BW. Arithmetic of subthreshold synaptic summation in a model CA1 pyramidal cell. Neuron. 2003;37:977–987.

99. Frey U, Morris R. Synaptic tagging: implications for late maintenance of hippocampal long-term potentiation. Trends Neurosci. 1998;21:181–188.

100. Smolen P, Baxter DA, Byrne JH. A model of the roles of essential kinases in the induction and expression of late long-term potentiation. Biophys J. 2006;90(8):2760–2775. doi:10.1529/biophysj.105.072470.

101. Tsokas P, Hsieh C, Yao Y, Lesburgueres E, Wallace EJ, Tcherepanov A, et al. Compensation for PKMZ in long-term potentiation and spatial long-term memory in mutant mice. ELife. 2016;5. doi:10.7554/eLife.14846.

102. Cerovic M, D’Isa R, Tonini R, Brambilla R. Molecular and cellular mechanisms of dopamine-mediated behavioral plasticity in the striatum. Neurobiol Learn Mem. 2013;105:63–80. doi:10.1016/j.nlm.2013.06.013.

103. Beninger RJ, Gerdjikov TV. Dopamine-Glutamate Interactions in Reward-Related Incentive Learning. In: Dopamine and Glutamate in Psychiatric Disorders. Totowa, NJ: Humana Press; 2005. p. 319–354. Available from: http://link.springer.com/10.1007/978-1-59259-852-6{_}14.

104. Yger M, Girault JA. DARPP-32, Jack of All Trades … Master of Which? Front Behav Neurosci. 2011;5(September):56. doi:10.3389/fnbeh.2011.00056.

105. Lindskog M, Kim M, Wikstriom MA, Blackwell KT, Kotaleski JH. Transient calcium and dopamine increase PKA activity and DARPP-32 phosphorylation. PLoS Comput Biol. 2006;2(9):e119. doi:10.1371/journal.pcbi.0020119.

106. Barbano PE, Spivak M, Flajolet M, Nairn AC, Greengard P, Greengard L. A mathematical tool for exploring the dynamics of biological networks. Proc Natl Acad Sci USA. 2007;104(49):19169–19174. doi:10.1073/pnas.0709955104.

107. Valjent E, Pascoli V, Svenningsson P, Paul S, Enslen H, Corvol JC, et al. Regulation of a protein phosphatase cascade allows convergent dopamine and glutamate signals to activate ERK in the striatum. Proc Natl Acad Sci USA. 2005;102(2):491–6. doi:10.1073/pnas.0408305102.

108. Devroyea C, Cathala A, Maitrea M, Piazzaa PV, Abrousa DN, Revesta JM, et al. Serotonin2C receptor stimulation inhibits cocaine-induced Fos expression and DARPP-32 phosphorylation in the rat striatum independently of dopamine outflow. Neuropharmacology. 2015;89:375–381. doi:http://dx.doi.org/10.1016/j.neuropharm.2014.10.016.

109. Hara M, Fukui R, Hieda E, Kuroiwa M, Bateup HS, Kano T, et al. Role of adrenoceptors in the regulation of dopamine/DARPP-32 signaling in neostriatal neurons. J Neurochem. 2010;113(4):1046–59. doi:10.1111/j.1471-4159.2010.06668.x.

110. D’Angelo E. The organization of plasticity in the cerebellar cortex: from synapses to control. Prog Brain Res. 2014;210:31–58. doi:10.1016/B978-0-444-63356-9.00002-9.

111. Kuroda S, Schweighofer N, Kawato M. Exploration of signal transduction pathways in cerebellar long-term depression by kinetic simulation. J Neurosci. 2001;21(15):5693–702.

112. Antunes G, De Schutter E. A stochastic signaling network mediates the probabilistic induction of cerebellar long-term depression. J Neurosci. 2012;32(27):9288–300. doi:10.1523/JNEUROSCI.5976-11.2012.

113. Hines ML, Morse T, Migliore M, Carnevale NT, Shepherd GM. ModelDB: A Database to Support Computational Neuroscience. J Comput Neurosci. 2004;17(1):7–11. doi:10.1023/B:JCNS.0000023869.22017.2e.

114. Chelliah V, Juty N, Ajmera I, Ali R, Dumousseau M, Glont M, et al. BioModels: ten-year anniversary. Nucleic Acids Res. 2015;43(Database issue):D542–548. doi:10.1093/nar/gku1181.

115. Sivakumaran S, Hariharaputran S, Mishra J, Bhalla US. The Database of Quantitative Cellular Signaling: management and analysis of chemical kinetic models of signaling networks. Bionformatics. 2003;19(3):408–15.

116. Lloyd CM, Lawson JR, Hunter PJ, Nielsen PF. The CellML Model Repository. Bionformatics. 2008;24(18):2122–3. doi:10.1093/bioinformatics/btn390.

117. Li C, Donizelli M, Rodriguez N, Dharuri H, Endler L, Chelliah V, et al. BioModels Database: An enhanced, curated and annotated resource for published quantitative kinetic models. BMC Syst Biol. 2010;4:92. doi:10.1186/1752-0509-4-92.

118. Le Novere N, Finney A, Hucka M, Bhalla US, Campagne F, Collado-Vides J, et al. Minimum information requested in the annotation of biochemical models (MIRIAM). Nat Biotechnol. 2005;23(12):1509–15. doi:10.1038/nbt1156.

119. Kiitter R. Postsynaptic integration of glutamatergic and dopaminergic signals in the striatum. Prog Neurobiol. 1994;44(2):163–196.

120. Hernandez AI, Blace N, Crary JF, Serrano PA, Leitges M, Libien JM, et al. Protein kinase MZ synthesis from a brain mRNA encoding an independent protein kinase CZ catalytic domain: implications for the molecular mechanism of memory. J Biol Chem. 2003;278(41):40305–40316. doi:10.1074/jbc.M307065200.

121. Maglott D, Ostell J, Pruitt KD, Tatusova T. Entrez Gene: gene-centered information at NCBI. Nucleic Acids Res. 2011;39(Database issue):D52–57. doi:10.1093/nar/gkq1237.

122. Whittaker V, Michaelson I, Kirkland RJA. The separation of synaptic vesicles from nerve-ending particles (synaptosomes). Biochem J. 1964;90(2):293.

123. Bai F, Weizmann FA. Synaptosome proteomics. In: Subcellular Proteomics. Springer; 2007. p. 77–98.

124. Vastagh C, Rodolosse A, Solymosi N, Liposits Z. Altered expression of genes encoding neurotransmitter receptors in GnRH neurons of proestrous mice. Front Cell Neurosci. 2016;10.

125. Silverman AJ, Hou-Yu A, Chen WP. Corticotropin-releasing factor synapses within the paraventricular nucleus of the hypothalamus. Neuroendocrinology. 1989;49(3):291–299.

126. Mystek P, Tworzydlo M, Dziedzicka-Wasylewska M, Polit A. New insights into the model of dopamine D1 receptor and G-proteins interactions. BBA-Mol Cell Res 2015;1853(3):594–603.

127. Ahn JH, Sung JY, McAvoy T, Nishi A, Janssens V, Goris J, et al. The B”/PR72 subunit mediates Ca^2+^-dependent dephosphorylation of DARPP-32 by protein phosphatase 2A. Proc Natl Acad Sci USA. 2007;104(23):9876–81. doi:10.1073/pnas.0703589104.

128. The Gene Ontology Consortium. Gene Ontology Consortium: going forward. Nucleic Acids Res. 2015;43(D1):D1049. doi:10.1093/nar/gku1179.

129. Fabregat A, Sidiropoulos K, Garapati P, Gillespie M, Hausmann K, Haw R, et al. The Reactome pathway Knowledgebase. Nucleic Acids Res. 2016;44(D1):D481. doi:10.1093/nar/gkv1351.

130. Kibbe WA, Arze C, Felix V, Mitraka E, Bolton E, Fu G, et al. Disease Ontology 2015 update: an expanded and updated database of human diseases for linking biomedical knowledge through disease data. Nucleic Acids Res. 2015;43(D1):D1071. doi:10.1093/nar/gku1011.

131. Jimeno-Yepes AJ, Sticco JC, Mork JG, Aronson AR. GeneRIF indexing: sentence selection based on machine learning. BMC Bioinformatics. 2013;14(1):171.

132. McKusick VA. Mendelian inheritance in man: a catalog of human genes and genetic disorders. vol. 1. JHU Press; 1998.

133. Amberger J, Bocchini CA, Scott AF, Hamosh A. McKusick’s online Mendelian inheritance in man (OMIM®). Nucleic Acids Res. 2009;37(suppl 1):D793–D796.

134. Chen Y, Cunningham F, Rios D, McLaren WM, Smith J, Pritchard B, et al. Ensembl variation resources. BMC Genomics. 2010;11(1):293.

135. He X, Simpson TI. statbio/topOnto: topOnto v1.0; 2017. Available from: https://doi.org/10.5281/zenodo.819735.

136. He X, Simpson TI. statbio/OntoSuite-Miner: OntoSuite-Miner v1.0; 2017. Available from: https://doi.org/10.5281/zenodo.819726.

137. Qi Z, Miller GW, Voit EO. Computational systems analysis of dopamine metabolism. PLoS ONE. 2008;3(6):e2444.

138. Sass MB, Lorenz AN, Green RL, Coleman RA. A pragmatic approach to biochemical systems theory applied to an α-synuclein-based model of Parkinson’s disease. J Neurosci Methods. 2009;178(2):366–377.

139. Harrison PJ, Weinberger DR. Schizophrenia genes, gene expression, and neuropathology: on the matter of their convergence. Mol Psychiatr. 2005;10(1):40.

140. Miannistoi PT, Kaakkola S. Catechol-O-methyltransferase (COMT): biochemistry, molecular biology, pharmacology, and clinical efficacy of the new selective COMT inhibitors. Pharmacol Rev. 1999;51(4):593–628.

141. Weinshilboum RM, Otterness DM, Szumlanski CL. Methylation pharmacogenetics: catechol O-methyltransferase, thiopurine methyltransferase, and histamine N-methyltransferase. Annu Rev Pharmacol. 1999;39(1):19–52.

142. De Araujo ME, Erhart G, Buck K, Muller-Holzner E, Hubalek M, Fiegl H, et al. Polymorphisms in the gene regions of the adaptor complex LAMTOR2/LAMTOR3 and their association with breast cancer risk. PLoS ONE. 2013;8(1):e53768.

143. Sterratt D, Graham B, Gillies A, Willshaw D. Principles of Computational Modelling in Neuroscience. Cambridge, UK: Cambridge University Press; 2011.

144. Hodgkin AL, Huxley AF. A quantitative description of membrane current and its application to conduction and excitation in nerve. J Physiol (Lond). 1952;117:500–544.

145. Gray KA, Seal RL, Tweedie S, Wright MW, Bruford EA. A review of the new HGNC gene family resource. Hum Genomics. 2016;10:6. doi:10.1186/s40246-016-0062-6.

146. Larson SD, Martone ME. NeuroLex.org: an online framework for neuroscience knowledge. Front Neuroinform. 2013;7:18. doi:10.3389/fninf.2013.00018.

147. Apweiler R, Attwood TK, Bairoch A, Bateman A, Birney E, Biswas M, et al. The InterPro database, an integrated documentation resource for protein families, domains and functional sites. Nucleic Acids Res. 2001;29(1):37–40. doi:10.1093/nar/29.1.37.

148. Sheridan RP, Venkataraghavan R. A systematic search for protein signature sequences. Proteins. 1992;14(1):16–28. doi:10.1002/prot.340140105.

149. Orengo CA, Bateman A, Uversky V, editors. Protein families: relating protein sequence, structure, and function. Wiley; 2014.

150. Rivals I, Personnaz L, Taing L, Potier MC. Enrichment or depletion of a GO category within a class of genes: which test? Bionformatics. 2007;23(4):401–7. doi:10.1093/bioinformatics/btl633.

151. Benjamini Y, Yekutieli D. The control of the false discovery rate in multiple testing under depencency. Ann Stat. 2001;29(4):1165–1188. doi:10.1214/aos/1013699998.

152. Alexa A, Rahnenfuihrer J, Lengauer T. Improved scoring of functional groups from gene expression data by decorrelating GO graph structure. Bionformatics. 2006;22(13):1600–1607. doi:10.1093/bioinformatics/btl140.

